# Comparative genomics of StAR-related lipid transfer (START) domains across wild and cultivated rice

**DOI:** 10.1101/2020.07.20.211607

**Authors:** Sanjeet Kumar Mahtha, Ravi Kiran Purama, Gitanjali Yadav

## Abstract

START proteins, encoded by a plant amplified family of evolutionary conserved genes, are known to bind lipids/sterols in mammals. In the plant kingdom, they have been reported in proteins involved in lipid binding, transport, signaling, as well as modulation of transcriptional activity, but there is very limited information on sub-functionalization or evolution of new function within the amplified gene family. Availability of ten fully annotated wild and cultivated rice varieties enables a comprehensive analysis of this family across the rice pangenome, across functional categories, as well as across START sub-families. The presence of START domains in all seven wild and three cultivated rice genomes suggests low dispensability and critical functional roles for this family, further supported by chromosomal mapping, duplication analysis and conservation of domain structure. Analysis of synteny highlights a preponderance of segmental and dispersed modes of duplication among STARTs, while transcriptomic investigation of the main cultivated variety *Oryza sativa var. japonica* reveals sub-functionalization among START genes family members in terms of preferential expression across various developmental stages and anatomical parts, especially flowering related parts. Importance of this family was further supported by Ka/Ks ratios confirming strong negative/purifying selection, implying that START family duplication is strongly constrained and evolutionarily stable.

## Introduction

The steroidogenic acute regulatory protein (StAR) related lipid transfer (START) domain was initially identified and named after mammalian StAR protein of 30 kDa, which binds to cholesterol [1]. START domains are evolutionarily conserved domains of approximately 200-210 amino acids [2] and play a crucial role in the transfer of lipids/sterols, lipid signaling and modulation of transcription activity [3,4]. Presence of START domains across evolutionarily distant organisms indicates a conserved mechanism for protein - lipid/sterol interaction through hydrophobic pockets [5]. Interestingly, START domains are abundant in plants and often associated with Homeodomain (HD) transcription factors, a feature unique to the plant kingdom [6]. For instance, 21 of the 90 homeodomain (HD) family members identified in Arabidopsis possess START domains along with putative leucine zippers [7]. Of these 21, five are from class III HD-ZIP subfamily and 16 1from class IV HD-ZIP subfamily [6,8].

The five genes from the class III HD-ZIP subfamily, namely PHB (Phabulosa), PHV (Phavoluta), REV (Revoluta), CAN (Corona) and ATHB8, have multiple and partially overlapping roles in development, including vasculature, organ polarity and embryonic patterning of the shoot meristem [9]. In contrast, several members of the class IV HD-ZIP subfamily have roles in layer specific cell differentiation. ATML1 (*Arabidopsis thaliana* Meristem Layer 1) and PDF2 (Protodermal Factor 2) have putative roles in epidermal differentiation [10,11]. GL2 (Glabra 2) is required for the differentiation of epidermal cells in the shoot [12], root [13] and seed [14]. ROC1 (Rice Outer Most Cell Specific Gene 1) of rice has similar function as ATML1, where its expression is limited to the outermost epidermal layer from the early embryogenesis [15]. OSTF1 (*Oryza sativa* Transcription Factor 1) also preferentially expressed in epidermis, and developmentally regulated during early embryogenesis [16].

Since HD START domain proteins act as transcription factors in plants, a major expectation is that START, when it binds to sterol, regulates gene expression similar to steroid hormone receptors from animals, this mechanism would allow cell differentiation to be linked with lipid metabolism in plants [3,17]. Plant START domains were shown to be required for transcription factor activity in class IV HD-ZIP protein ‘GL2’ in Arabidopsis, and they were also found to have ligand-binding modules, similar to mammalian START domains [18]. Activated expression of HDG11 START domain confers drought tolerance with reduced stomatal density and improved root system in Arabidopsis [19].

Although START domains are amplified in plants, and appear to have diverse functions, a thorough knowledge of mechanism of amplification and gene duplication in this family is lacking. With availability of many varietal genomes, and huge genotypic variation ranging from diploids to polyploids, Oryza has a long history of use as a model monocot food crop. Furthermore, with a wide evolutionary history that spans more than 15 million years, Oryza is an ideal prototype for such a study [20–22]. Rice also has a major social significance, consumed by half the global population with an estimated 20% of human dietary calories that are met only by the domesticated Asian rice variety, thereby making it a target for improvement towards addressing the food security issue of a growing world population under a changing climate.

In this work, we focus on ten diploid Oryza species, including three cultivated varieties having AA genotype, *O. sativa var. indica*, *O. sativa var. japonica* (Asian cultivated variety), along with *O. glaberima (african cultivated variety).* Five of the seven wild Oryza species included in this analysis have AA genotype, namely *O. rufipogon, O. nivara, O. barthii, O. glumaepatula*, and *O. meridionalis,* while two others have BB and FF genotypes, *O. punctata* and *O. brachyantha* respectively. Regardless of genotypes, all ten species have enormous repeats, varying from one fourth to half the genome size [22]. In general, repeat regions are accumulated with increasing evolutionary order from early-evolved wild relatives such as *O. brachyantha* and *O. meridionalis* (approx 27-29%) to the recent cultivated varieties *O. sativa var. japonica* and *indica* (approx 40-50%). *O. punctata* is an exception, despite being an early evolved wild species, has half of its genome containing repeats; resulting in a huge repertoire of synteny within the Oryza genome, varying from 90% to 97% [22]. In addition there is an gene flow among AA type Oryza genomes, which needs to be thoroughly investigated to understand the specific changes that occurred in the gene families [21,22]. The expanded gene family of START domains can be single or multi-domain [6,23], and has been reported to associate with several other domains such as homeodomain, MEKHLA, and PH domains, known for their involvement in transcription regulation, sensing and signaling, respectively[3,6,17,24].

In this paper, we aim to provide a comprehensive comparative genomic analysis of START genes across the ten Oryza genomes, investigated all the way from identification and classification to sequence homology, genome-wide mapping, and duplication analysis of START genes. Available transcriptomic data for *Oryza sativa var. japonica* was investigated to understand co-expression patterns for potential sub- or neo-functionalization among these genes. Genome wide identification revealed a total of 249 START genes taking all ten rice species together, and showed that the gene family size for START genes varies from 22 to 28. Domain structure analysis confirmed presence of additional functional domains associated with STARTs such as HDs, MEKHLA, PH, and DUF1336 and classified the START proteins into total eight unique combinations based on associated domain patterns. Phylogenetics revealed the extent of divergence amongst START proteins and we find distinct clusters based on above mentioned domain structure patterns. The genome-wide mapping showed that these genes are distributed among 11 chromosomes out of 12 in most of the cultivated and wild rice species. Gene duplication studies indicate that, START genes preferred segmental and dispersed modes of duplication for gene expansion under natural selection. Hierarchical clustering of transcriptome data revealed many duplicated gene pairs have similar expression patterns across developmental stages and anatomy. In summary, this is a comparative genomics of START genes across wild and cultivated rice, and enhances our understanding about the mechanism of START gene amplification in plants.

## Materials and Methods

### Data collection

The complete genomic sequences, protein sequences, and annotation information of nine species of oryza, including seven wild varieties *Oryza brachyantha (Obra_w_), Oryza punctata (Opun_w_), Oryza meridionalis (Omer_w_), Oryza glumaepatula (Oglu_w_), Oryza barthii (Obar_w_), Oryza nivara (Oniv_w_), Oryza rufipogon (Orup_w_)* along with two cultivated varieties *Oryza glaberrima (Ogla_c_),* and *Oryza sativa var. indica (Oind_c_)*, were downloaded from Ensembl [25]. In addition, similar data for the main cultivated variety, *Oryza sativa var. japonica (Ojap_c_)* was downloaded from the Phytozome v12 having latest updated version of sequences [26]. Throughout this work, these ten species are referred to in subscripted ***Oabc*_*x*_** format where *abc* represents first three letters of the species/subspecies name, while the subscript ‘x’ is *c or w*, representing cultivated or wild nature respectively.

### Identification & validation of START proteins

Previously reviewed and characterized sequences of 109 START domain-containing proteins were collected from Interpro consortium [27]. The START regions in these proteins were extracted based on annotated border residues, and sequence redundancy was removed at cut off 95% using CD-hit [28] The resulting 84 sequences were used to construct a profile hidden markov model with HMMER 3.2.1 (http://hmmer.org/) [29,30]. The profile was run against all ten Oryza proteomes and short hits (sequence length < 100 residues) were discarded, followed by removal of redundancy, performed by filtering out all but the longest peptide for each protein. The validation of identified hits as START family proteins was performed using Conserved Domain Database (CDD) [31].

### Domain structure analysis of START domain containing proteins

The putative START domain containing proteins identified as described above, were subjected to domain structural pattern analysis to ascertain additional domains associated with START. Domain structure analysis was carried out using a web based Batch CD-search Tool, selecting Conserved Domain Database (CDD) [32]. CDD includes curated data from NCBI (National Center for Biotechnology Information) [33] SMART (Simple Modular Architecture Research Tool) [34] Pfam (protein families) database [35], PRK (**PR**otein **K**(c)lusters) [36], COG (**C**lusters of **O**rthologous **G**roups of proteins) [37], and TIGRFAMs (**T**he **I**nstitute for **G**enomic **R**esearch’s database of **p**rotein **fam**ilies) [38]. The additional associated domains, as identified in this step were used to classify rice START domains into various domain structural classes. Besides this trans-membrane helical segments associated with START domains were predicted using TMHMM Server v. 2.0 [39].

### Gene structure analysis of START coding genes

Gene structure analysis was carried out to understand the exon-intron patterns for different classes of START encoding genes among ten rice species. The corresponding Gene and CoDing Sequence (CDS) of the START encoding proteins were used as input for gene structure analysis. Gene structure was visualized using Gene structure Display Server (GDSD) [40].

### Genome-wide mapping and Identification of Homologous and Orthologous START coding genes amongst ten Oryza species

In order to map the START coding genes onto rice chromosomes, gene location data was extracted from the respective GFF annotation files (General feature format), and karyotype information was extracted based on chromosomal length. Chromosomal visualization of genes in all ten rice species was done using circos [41], colored by structural class. Orthologous START genes in nine Oryza species were identified in reference to *Ojap_c_* by local protein BLAST, based on maximum identity and similarity.

### Phylogenetic analysis of different structural classes of START domain containing proteins

Phylogenetic analysis was carried out for different structural classes of START proteins across all ten species, to explore intra- and inter-species divergence. All 249 full-length START proteins in the ten Oryza genomes and 35 sequences from *Arabidopsis thaliana* were included in the phylogenetic study. The available gene symbols are used in case of *O. sativa var. japonica* and *Arabidopsis thaliana.* Multiple sequence alignment was performed by Clustal omega at default parameters [42]. Aligned sequences were used for phylogenetic tree construction. The tree was generated through PHYLIP v3.2 [43] using neighbor-joining method at bootstrap value of 1000 and the tree was visualized using Figtree v1.4.2 [44] (http://tree.bio.ed.ac.uk/software/figtree/).

### Gene Duplications, collinearity and nucleotide substitution rates

The MCScanX software package [45] was used to identify various duplication modes for START genes among Oryza species. This program works on the all-vs-all BLASTp results and this was performed for all ten rice proteomes [46]. The results were fed into duplicate gene classifier, a module of MCScanX, to detect dispersed, proximal, tandem and/or segmental duplications. Unduplicated genes (that occur only once in the genome) were classified as singletons. Collinear blocks for all proteins within individual genomes were generated by MCScanX module (grey colour links). START gene homologues within collinear blocks were highlighted using the previously described domain structure class colours. MCScanX transposed [47] was used to find the newly trans-located START homologues from their original ancestral locations to a novel locus in *Ojap_c_*. The START gene homologues obtained from the interspecies BLASTp between *Ojap_c_* and the other nine Oryza genomes were analysed for non-synonymous (Ka) and synonymous (Ks) substitution rates by KaKs calculator 2.0 [48] for *Ojap_c,_* all START homologues in collinear blocks were extracted from the collinearity files generated in MCScanX-transposed runs.

### Transcriptome analysis and hierarchical clustering

Gene expression levels of 28 START genes in the major globally cultivated rice variety *Ojap_c_* were investigated using RNA-seq data ‘Os_mRNAseq_Rice_GL-0’ (MSU v7.0) on the Genevestigator platform [49]. The conditional search tool was used to analyse gene expression across nine developmental stages and 13 anatomical parts, and their log transformed values were further arranged in hierarchical clustering groups based on Pearson correlation coefficients of START genes by selecting optimal leaf ordering for both, developmental stages and anatomical parts. The heatmaps were generated using Mev_v4.8[50]

## Results

### Identification of START genes amongst wild and cultivated varieties of Rice

The HMM profiles, constructed based on known sequences, were used to perform the HMM search against ten *Oryza* proteomes (listed in Table 1) and only those hits were retained that matched the minimum length criteria, and were validated for presence of START, as described in Methods. This led to the identification of 360 START proteins (including protein isoforms), coded by 249 gene transcripts across the ten species of rice. In order to remove redundancy, only the single longest protein coded by each set of gene transcripts was retained for downstream analysis. START coding genes were found to vary from 22 to 28 in these Oryza species as shown in Table 1.

**Table 1.**
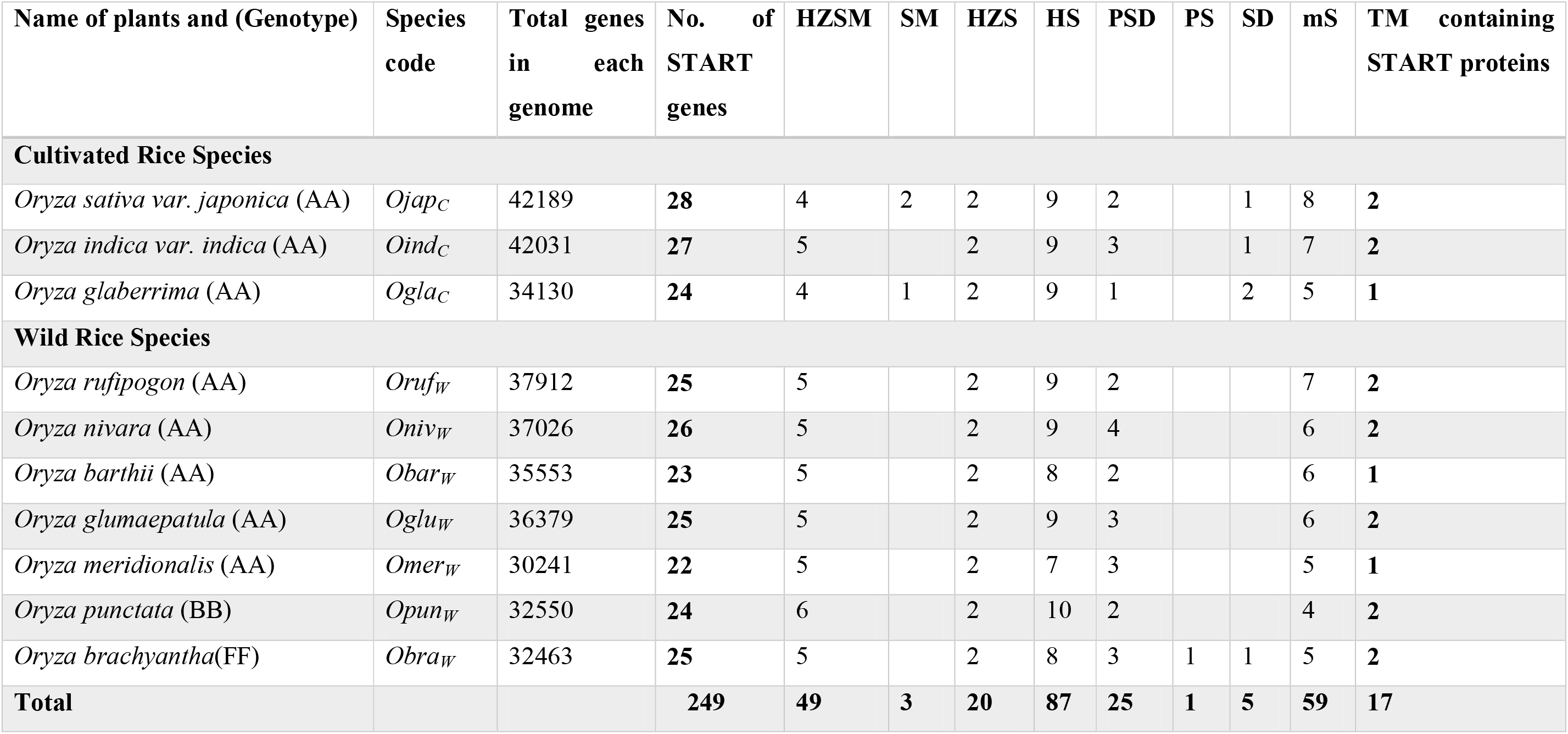
Identification and Domain structure analysis of START proteins across cultivated and wild rice species. (**HZSM**: HD bZIP START MEKHLA, **SM**: START MEKHLA, **HZS**: HD bZIP START, **HS**: HD START, **PSD**: PH START DUF1336, **PS**: PH START, **SD**: START DUF1336, **mS**: minimal START, **TM**: Transmembrane segments). The species are first categorized based on their cultivated and wild nature and then ranked in the order of evolution, from recent to oldest evolved.

| Name of plants and (Genotype) | Species code | Total genes in each genome | No. of START genes | HZSM | SM | HZS | HS | PSD | PS | SD | mS | TM containing START proteins |
| --- | --- | --- | --- | --- | --- | --- | --- | --- | --- | --- | --- | --- |
| **Cultivated Rice Species** |  |  |  |  |  |  |  |  |  |  |  |  |
| *Oryza sativa* var. *japonica* (AA) | *Ojap<sub>C</sub>* | 42189 | **28** | 4 | 2 | 2 | 9 | 2 |  | 1 | 8 | **2** |
| *Oryza indica* var. *indica* (AA) | *Oind<sub>C</sub>* | 42031 | **27** | 5 |  | 2 | 9 | 3 |  | 1 | 7 | **2** |
| *Oryza glaberrima* (AA) | *Ogla<sub>C</sub>* | 34130 | **24** | 4 | 1 | 2 | 9 | 1 |  | 2 | 5 | **1** |
| **Wild Rice Species** |  |  |  |  |  |  |  |  |  |  |  |  |
| *Oryza rufipogon* (AA) | *Oruf<sub>W</sub>* | 37912 | **25** | 5 |  | 2 | 9 | 2 |  |  | 7 | **2** |
| *Oryza nivara* (AA) | *Oniv<sub>W</sub>* | 37026 | **26** | 5 |  | 2 | 9 | 4 |  |  | 6 | **2** |
| *Oryza barthii* (AA) | *Obar<sub>W</sub>* | 35553 | **23** | 5 |  | 2 | 8 | 2 |  |  | 6 | **1** |
| *Oryza glumaepatula* (AA) | *Oglu<sub>W</sub>* | 36379 | **25** | 5 |  | 2 | 9 | 3 |  |  | 6 | **2** |
| *Oryza meridionalis* (AA) | *Omer<sub>W</sub>* | 30241 | **22** | 5 |  | 2 | 7 | 3 |  |  | 5 | **1** |
| *Oryza punctata* (BB) | *Opun<sub>W</sub>* | 32550 | **24** | 6 |  | 2 | 10 | 2 |  |  | 4 | **2** |
| *Oryza brachyantha*(FF) | *Obra<sub>W</sub>* | 32463 | **25** | 5 |  | 2 | 8 | 3 | 1 | 1 | 5 | **2** |
| **Total** |  |  | **249** | **49** | **3** | **20** | **87** | **25** | **1** | **5** | **59** | **17** |

As can be seen from this Table, the most widely cultivated varieties *Ojap_c_*, and *Oind_c_* possess the largest numbers of START coding genes, when compared to early evolved African cultivated species *Ogla_c_* and the seven wild species. The oldest AA variety *Omer_w_* has 22 START coding genes, lowest among all, although START numbers do not vary greatly between species, and their numbers are proportional to genome size in most cases. Among the wild varieties, the earliest evolved *Obra_w_* has the same number of START genes as the most recently evolved wild variety *Oruf_w_*. The breakup of these START domains, in terms of potential functions based on domain combinations, is explored further in the next section, but these general numbers suggest that the increase in START domains among cultivated rices, reflects a role of STARTs in stress. For accession IDs of START coding genes found in all ten species, along with protein and domain information, see Supplementary Table S1.

### Domain structure analysis (DSA) and classification of START proteins

START proteins have been known to exist both as minimal START domains, as well as in association with other functional domains, and this domain organization has been used as a criterion for their classification along with information on specific ligands, which they bind [6,23]. We explored the domain structure of all 249 identified START domains, and classified them into eight groups, as shown in Table 1, depending on the combinatorial patterns of STARTs with additional domains such as HD (Homeodomain), bZIP (Basic Leucine Zipper Domain), MEKHLA domains, PH (Pleckstrin homology domain) and DUF1336 (Domain of unknown function). Six of these eight groups have been reported earlier, including (a) mS i.e. minimal START lacking any additional domains, (b) HS (having HD), (c) HZS (containing HD and bZIP), (d) HZSM (having HD, bZIP and MEKHLA), (e) PSD (with PH and DUF1336), and (f) SD (START with DUF1336), while two new combinations (not reported earlier) were also seen, namely (g) SM (i.e START with MEKHLA) and (h) PS (START with PH). Interestingly, these last two combinations are the only ones that are either completely absent from the cultivated varieties (as in case of PS), or completely absent from wild varieties (as in case of SM).

As can be seen in Table 1, almost 24% of rice START domains belong to minimal START (i.e lacking any additional domains), while homeodomains constitute the largest category of domains co-occurring with STARTs. The recently evolved cultivated rices (*Ojap_c_ or Oind_c_)* have higher number of minimal STARTs compared to early evolved wild species *(Obra_w_ or Opun_w_)*. We have previously shown that the HD associated with STARTs in plants has unique roles in plant transcription [18], and this seems to be ancient feature, since all wild rice species also have the homeodomains. The HDs are always found in association with a leucine zipper in class III and class IV HD-zip family of plant START proteins. Over 60% of the 249 identified domains in rice have these homeodomains in combination with leucine zippers, which in turn, can be of two types; (a) class III HD ZIP START domains with a universally conserved basic leucine zipper known as bZIP, and (b) class IV HD ZIP STARTs, with a plant exclusive leucine zipper, known as ZLZ [6,8]. Another domain, MEKHLA, often seen associated with the class III HD bZIP START proteins [24], is completely missing from the START domains in all the wild rice species (SM family), as can be seen from Table 1. Our DSA methodology is based on Conserved domain Database [31] which does not recognize the ZLZ, hence we use the term ‘HD-START’ for class IV type proteins throughout this study.

Interestingly, the difference between domain structure of wild and cultivated rice does not appear to arise from the homeodomain containing STARTs, all of which occur in large numbers and with moderate uniformity across all rices (See Table 1). Apart from the HD containing START domains, the other two major domains that co-occur with STARTs are the pleckstrin homology at the N-terminus, and DUF 1336 domains at the C-terminus. These form unusual combinations, two of which have been observed for the first time in this work, as mentioned earlier, and are starkly distinct between wild and cultivated rices; the dual combinations of START DUF1336 (five), START MEKHLA (three), and PH START (one). Infact, a START domain in combination with the pleckstrin homology alone, has only been observed in the earliest evolved wild rice namely, *O.bra_w_*. Similarly, very few domains show the combination of START domain alone with DUF1336, but the triple combination (PSD category) is seen frequently (35% of non-HD START combinations) across all rices, suggesting that these three domains are more effective in combination, rather than alone. PH domains are well known for intracellular signaling or as constituents of the cytoskeleton proteins. This domain also binds with phosphatidylinositol within biological membranes, thus playing roles in membrane recruitment, sub cellular targeting or enabling interactions with other components of the signal transduction pathways [51,52]. Intriguingly, another connection to this role is evident in minimal STARTs, 30% of which were found to have transmembrane (TM) segments (17 in all; 11 with two TM segments and six with single TM), that show a huge similarity to a specific class of mammalian STARTs, namely the Phosphatidylcholine transfer proteins (STARD2/PCTP) that preferentially bind to Phosphatidylcholine [53]. That PH domains are present singly with START domains in the earliest known rice, and not elsewhere, as well as the presence of TM segments, but only in minimal STARTs, suggests that initiation of association with other domains began with membrane interfaces, and the addition of other, newer domains, may have been critical to evolution of START functional diversity. The illustrative image for domain organization of 28 START proteins from *Ojap_c_* is given in Figure 1. The detailed domain structure analysis report of 249 START proteins along with the domain sizes and positions is provided in Supplementary Table S1.

**Figure 1.**
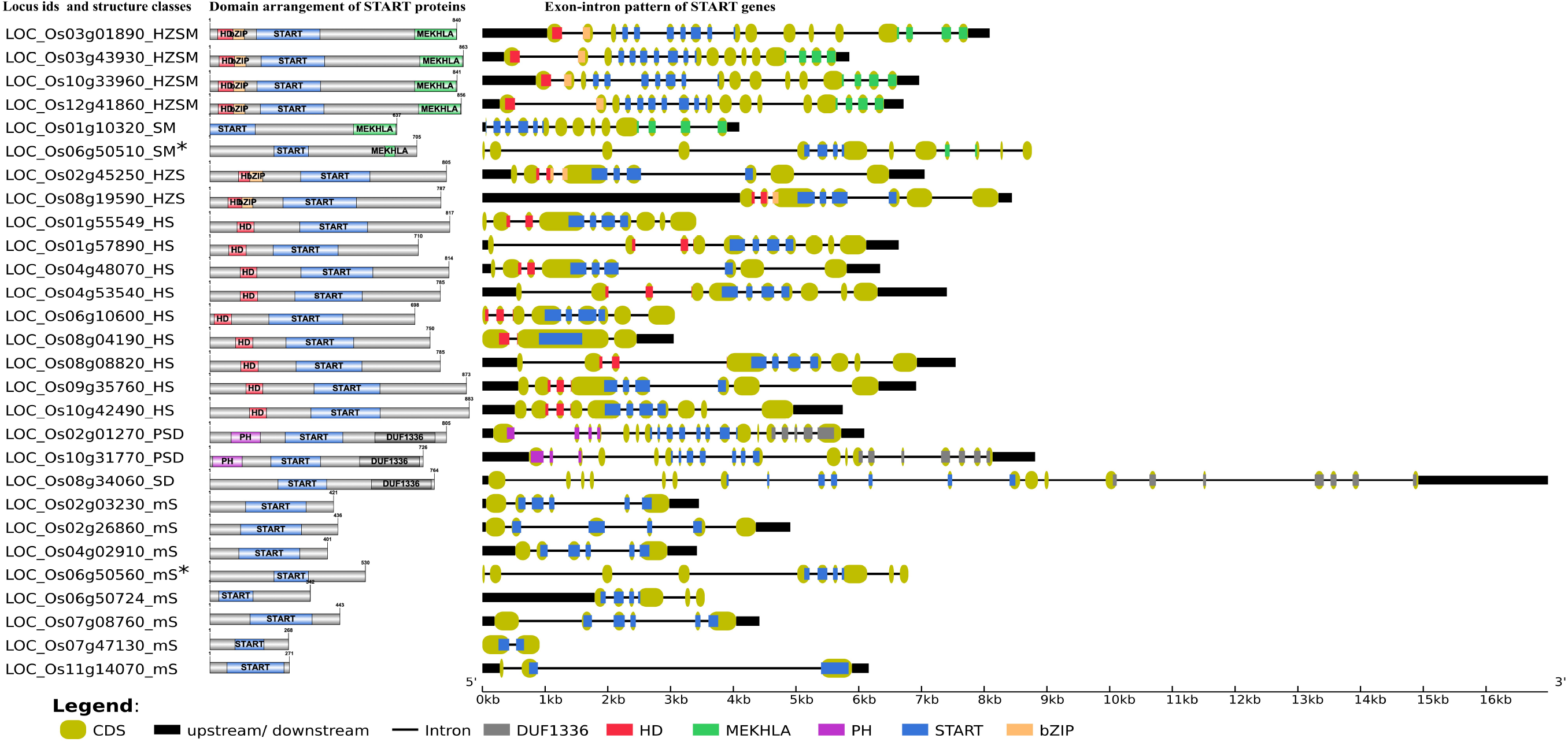
Schematic overview of domain arrangement of START proteins generated using IBS v.1.0 [54] and gene structure of all 28 STARTs genes in *Oryza sativa var. japonica.* Index shows exonic colors for homeodomain (HD), homeodomain associated basic Leucine zipper domain (HD bZIP), START domain, PH domain, Domain of Unidentified Function (DUF1336), and MEKHLA. The exon portions that code the protein inter-domain regions (grey colored regions in Domain arrangement) are depicted in mint shades at gene structure. (*Gene pairs are proximally duplicated and shows higher resemblance in gene structure).

### Gene structure analysis (GSA) of START coding genes

GSA for all 249 START domains was performed as described in methods, and results are depicted in Table 2, listing exon numbers for each functional domain within and between the eight START categories described in the previous section. The gene structure analysis for main cultivated variety *Ojap_c_* along with its full-length protein domains pattern is depicted in Figure 1, whereas full gene structure maps and completes exon-intron details of all START domains in all ten rices are provided as supplementary data (Supplementary Figures S1A-J, and Supplementary Tables S1 & S2). In general, gene sizes vary between .5 kb to 32 kb, with some of the minimal STARTs in the oldest wild rice *Obra_w_* encoded by a single exon, while few PSD class STARTS in wild rices have upto 34 exons. The overall pattern (See Table 2) is that exon numbers for the START region itself are quite conserved within specific structural classes, and the variability between wild and cultivated rice stems from exon numbers of the associated domains in these proteins. Exon numbers are also highly variable across the eight classes of START domains, with greatest variability reflected in the minimal STARTs which contain very large intronic regions and long lengths of upstream and downstream UTRs (up to 17Kb), suggesting a potential for addition of new domains, exon creation and alternate splicing. Figure 1 reveals similarities between gene structure of the newly observed SM class of STARTs with other categories; GSA of one of the SM genes is almost identical with HZSM members, after losing the HZ fragment, whereas the other SM gene has a GSA identical to a member of the minimal STARTs (Figure 1), suggesting gain of function. They are also proximally duplicated where SM classes showed higher expression in both anatomical part and development stages while mS classes are poorly expressed (see section on gene expression). Both pairs of genes are on the same chromosomes, adding support to these hypotheses, as discussed in the subsequent sections on chromosomal mapping and phylogenetics.

**Table 2.**
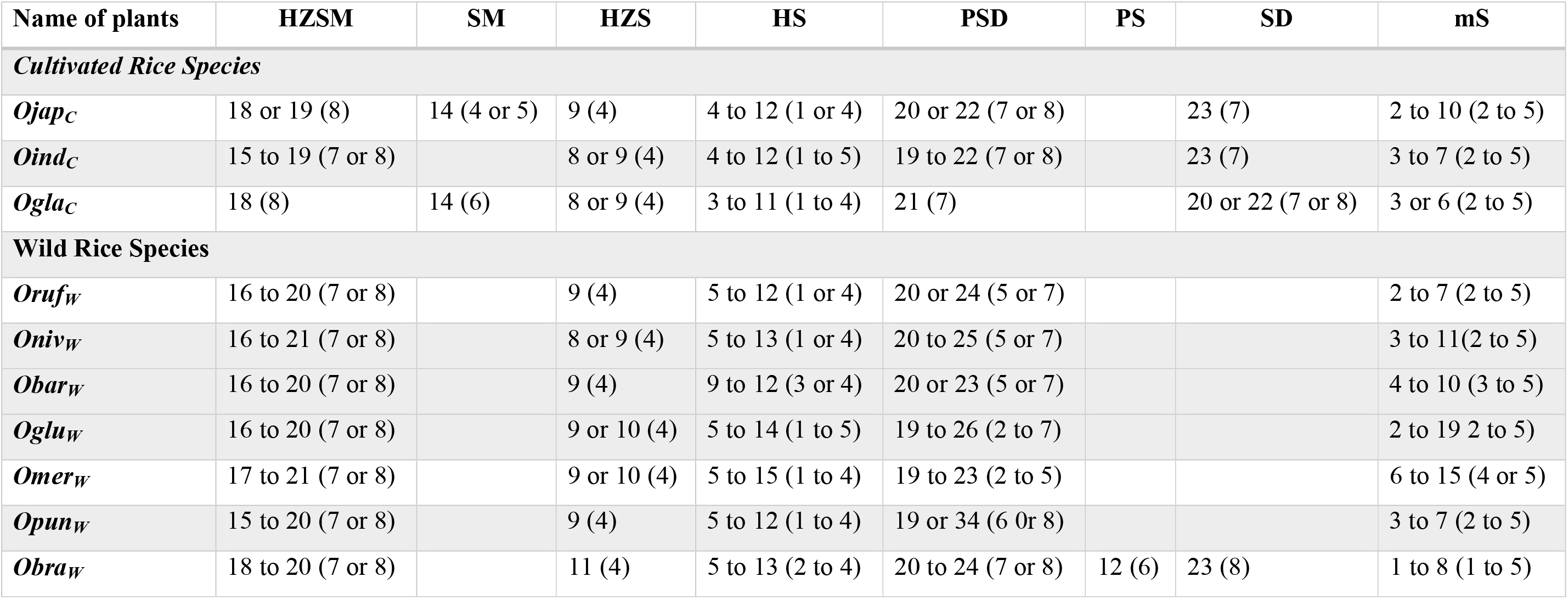
Gene structure analysis: Number of exons involve in coding full-length START proteins for different structural classes amongst ten cultivated and wild Oryza species. The values given in parentheses are the number of exons that code for START domains regions alone. Acronyms/Codes same as Table 1

Apart from the above mentioned differences in the number of exons, START coding genes also vary in intron length and Untranslated Regions (UTR) regions across the cultivated and wild rices. There is still much that is unknown about flanking untranslated regions at both the terminals of mRNA in the form of 3’ UTR and 5’ UTR, and although UTR regions have often been implicated in regulatory aspects of gene expression, they need to be investigated further. Almost one third of all START coding genes reveal long sections of 3’ UTR and 5’ UTR (ranging from few nucleotides upto 17 kb), but the African and Asian cultivated rices (*Oind_c_* and *Ogla_c_*) appear to completely lack these at both termini. Clearly, cultivated rices vary by ancestors, and this is reflected in their inherent genetic diversity, as observed between *Oind_c_ and Ojap_c_*. As can be seen in Supplementary Figure S1A-J and Supplementary Table S2, UTR lengths were observed to be very long in minimal START genes, along with very long intron lengths, both features suggesting the potential for evolution via introduction of new function. Among the various classes of START genes, HZSM shows the shortest exon and introns regions while PSD and SD classes show distinctive combination of several exons and longer introns, aside from long stretches of UTR regions. Most classes have exons flanked by long introns but the HZSM and PSD have exons flanked by short introns. Cultivated rices having fewer cases of long flanking introns, further emphasize the greater genetic diversity in wild rices and scope for exon creation, alternate splicing and addition of functional features.

### Ortholog analysis and chromosomal distribution

The putative START coding genes were mapped on to chromosomes based on their gene location and karyotype information. Figure 2 depicts this for all ten *Oryza* genomes and it is clear that despite variation in numbers, START genes show positional and structural conservation on the corresponding chromosomes, with slight variations in some genes reflecting syntenic block shuffling, which may in turn be due to (a) fragment rearrangements among chromosomes during speciation events, (b) isolated gene relocation events due to the homologues recombination or viral or transposon based gene relocation mechanisms.

**Figure 2.**
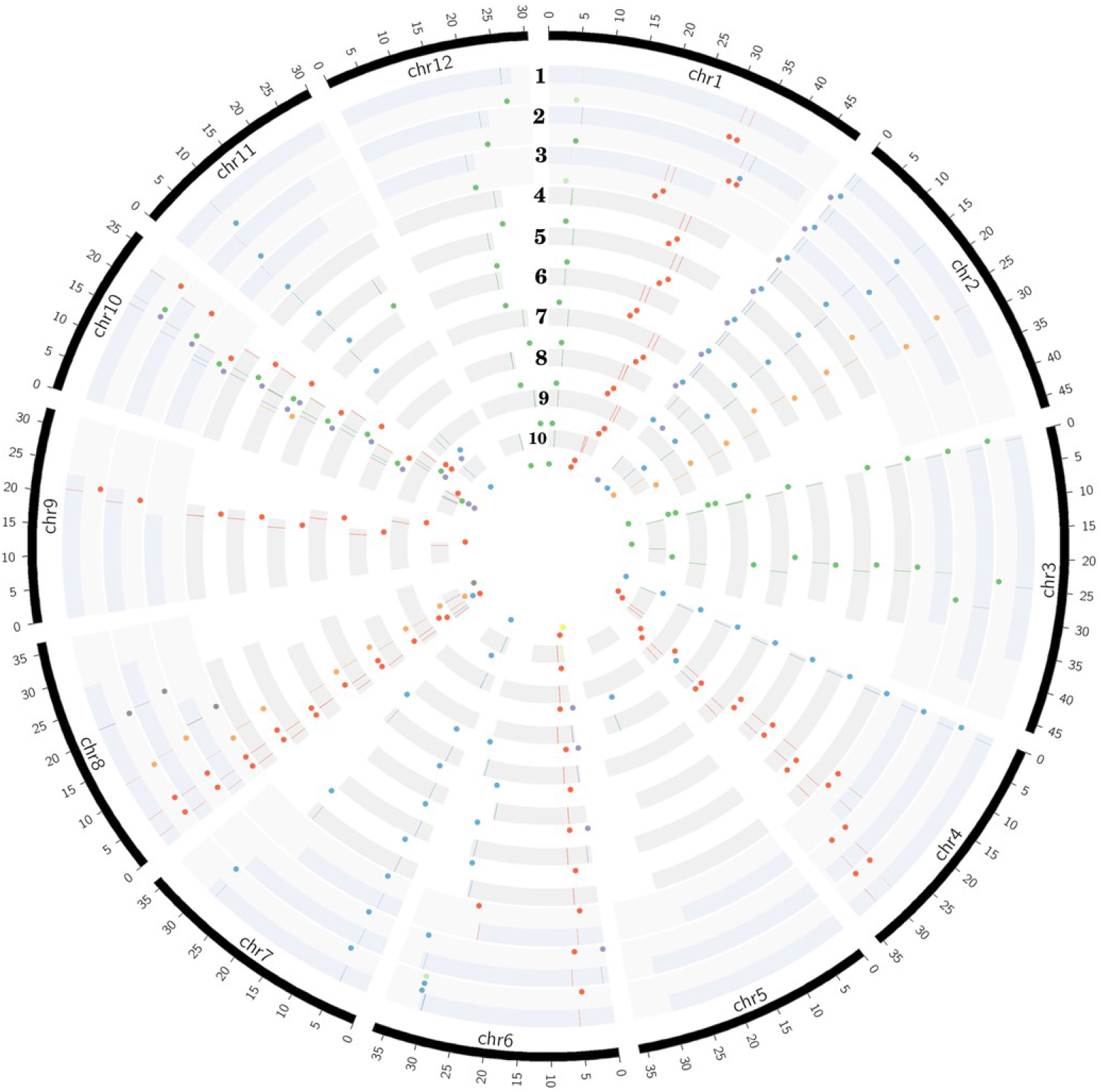
Distribution of all eight START categories (colored dots shown internal to chromosomes in circular ribbons and lines on chromosomes) across ten Oryza genomes (each ribbon representing a chromosome in circular form). The ten rices are arranged in order of Cultivated (1-3) and wild (4-10), and represented in evolutionary order from recently evolved to the oldest: 1- *Oryza sativa var. japonica*, 2- *Oryza sativa var. Indica,* 3- *Oryza glaberrima,* 4- *Oryza rufipogon,* 5- *Oryza nivara, 6- Oryza barthii,* 7- *Oryza glumaepatula*, 8- *Oryza meridionalis*, 9-*Oryza punctata* and 10- *Oryza brachyantha*. START types represented by circular glyphs/dots: Blue - Minimal START; Grey - SD; very light green - SM; yellow - PS; purple - PSD; red - HS; light orange - HZS; and green - HZSM.

START genes are distributed among 11 chromosomes (out of 12) across all wild and cultivated rices species (Figure 2), with the highest numbers mapping to chromosomes 8 and 10, while chromosome 5 is devoid of any START genes (with a single exception of one START gene in early evolved *Omer_w_*). The HZSM class of START genes is predominantly located on chromosome 1, 3, 10 and 12. Surprisingly, there are two HZS gene orthologues unequivocally present, one each on chromosome 2 and 8 except in *Oniv_w_ where* one HZS gene was seen on chromosome 10 instead of chromosome 8. It may be recalled that SM, a special class of START that was not seen in any of the wild rices, occurs on chromosome 1 amongst cultivated rices *Ojap_c_* and *Ogla_c_* and shares homology with HZSM on the same chromosome in eight other rices, suggesting a possible loss of HZ fragment of some members of the HZSM class. In contrast, the other SM gene on chromosome 6 of *Ojap_c_* showed homology with minimal START on chromosome 6 of *Oruf_w_*, *Oniv_w_*, *Obar_w_*, and *Oglu_w_* and is possibly an example of gain of function. These observations match the pairwise gene structure analysis patterns observed in the previous section, and are further supported by the corresponding pairs of genes being orthologous as shown in Table 3. This table lists the orthologous genes in all ten rices, using the recently evolved cultivated variety *Ojap_c_* as reference for the other nine rice genomes, as described in methods. Interestingly, this table also shows that PS, (a special class of STARTs, seen only in the oldest wild rice *Obra_w_*) is orthologous to a member of the PSD class in the cultivated varieties. In contrast, the members of the SD class, observed only in cultivated Oryza species, and the oldest wild rice, are similar to each other, but do not have any orthologs in other genomes of rice, not even in the immediate ancestors of the three cultivated varieties. Overall, the findings of this section support the idea of specialized functional roles for each of the eight START classes in plants, and we further explore this aspect in later sections.

**Table 3.**
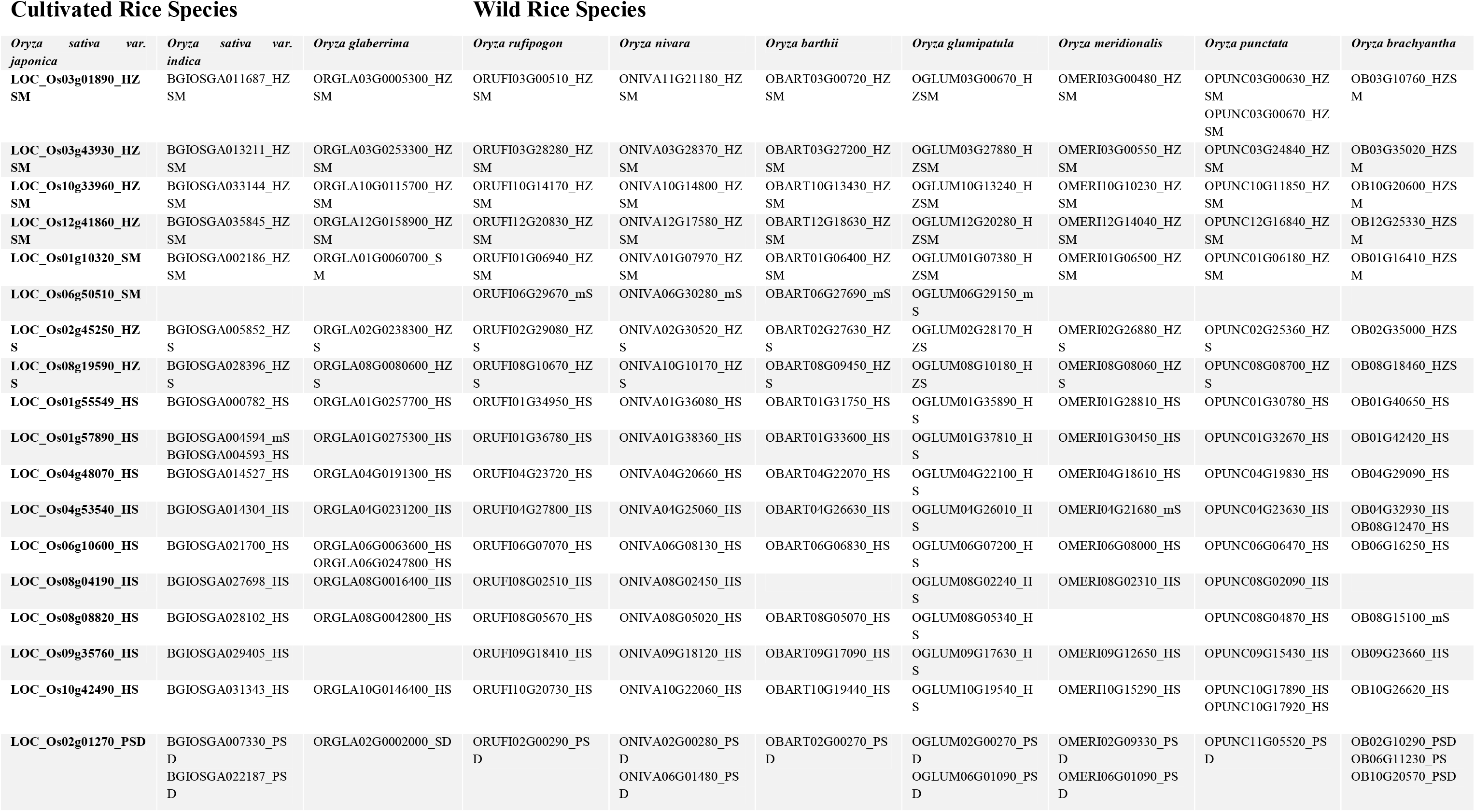
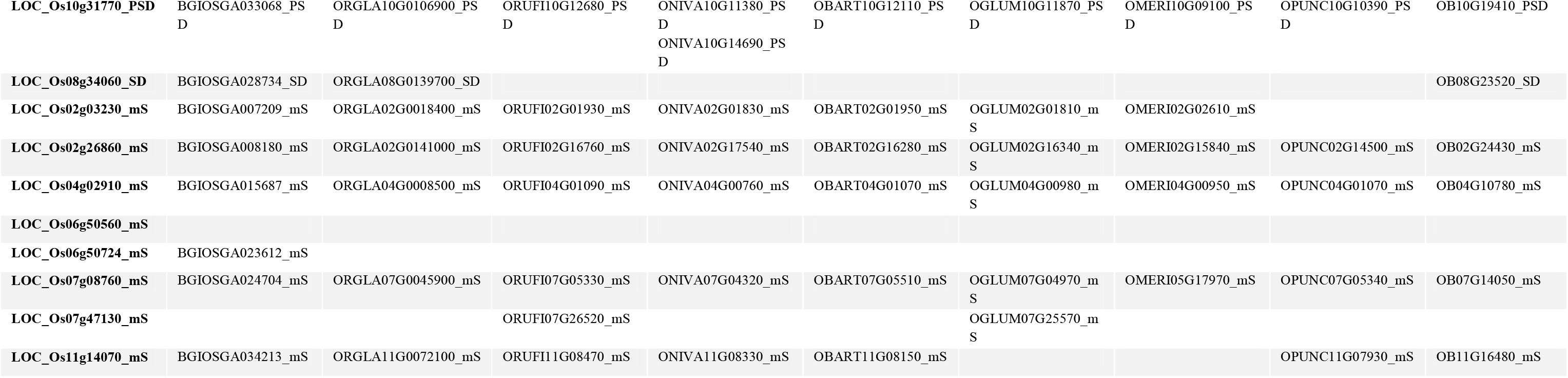
START genes in *Oryza sativa var. japonica* and their best Orthologs amongst other 9 rice species.

### Phylogenetic analysis

The 249 START protein sequences from all ten Oryza species and 35 reference sequences from model plant *Arabidopsis thaliana* were used to construct a phylogenetic dendrogram as described in methods, and this led to grouping of genes having closely related evolutionary patterns as shown in Figure 3. The phylogenetic tree showed distinct clusters for all major structural classes of START domains, which suggests conservation amongst different structural classes of START proteins in terms of their sequences. Few of the minimal STARTs were distributed among different clusters that might be due to their vast differences in their sequence lengths. As shown in Figure 3, The HZSM represented in green forms a single distinct cluster, while HZS and HS represented in light orange and red, respectively formed a single cluster, as expected with an overlap between these two sub classes. The two minor classes *i.e.* PS (shown in olive) and SD (shown in grey) formed sub-cluster together with PSD (shown in purple). The minimal START proteins represented in blue forms a single large cluster, but some of them are distributed among other structural classes, which might be due to vast differences in their sequence lengths. The three SM class (represented in dark green) falls alongside of HZSM and minimal START. As expected, all three cultivated rices were observed to lie adjacent to each other or in the same branch as their immediate wild ancestor.

**Figure 3.**
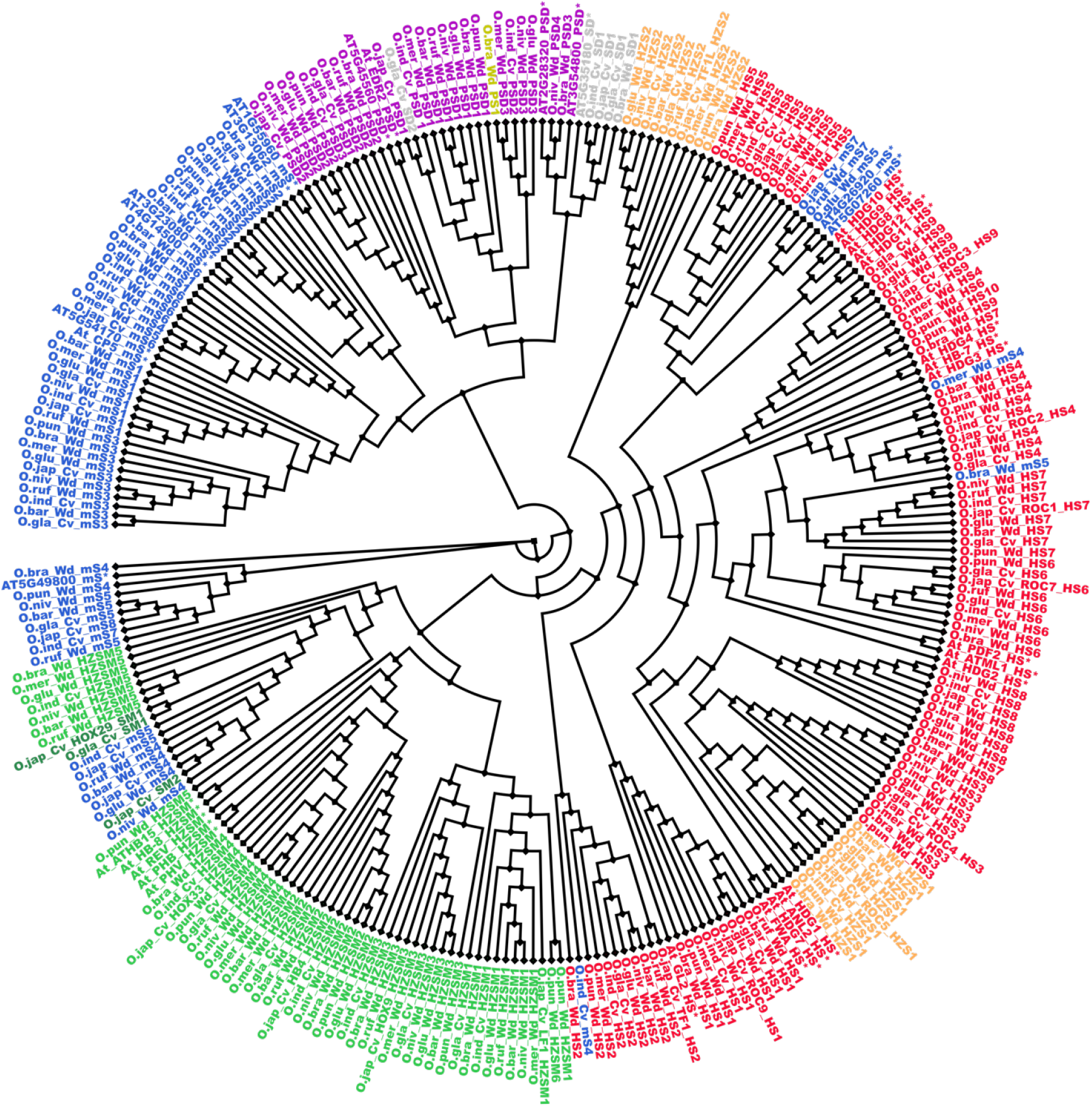
Cladogram for START proteins from all ten Oryza genomes along with *Arabidopsis thaliana* (*). Color codes are same as earlier figures: Red: HD START (HS); light orange: HD bZIP START (HZS); green: HD bZIP START MEKHLA (HZSM); dark green: START MEKHLA (SM); purple: PH START DUF1336 (PSD); yellow: PH START (PS); grey: START DUF1336(SD); blue for minimal START. Phylogeny codes for each locus ID are based on the orthologs analysis with reference to *Osj_c_* as described previously (See supplementary Table S1).

Previous studies suggested that Class III HD-ZIP proteins are evolutionarily conserved [8]. In this study, although this family formed a single cluster there were few intervening minimal STARTs were found. The unusual START type ‘START MEKHLA’ also merged with this cluster, which shows evolutionary relatedness, despite the lack of HD-ZIP region these proteins to class III HD-ZIP proteins. The HD bZIP START and HD START shared the high similarity between the two and which causes overlapping of clusters. Similarly PS, PSD, and SD are forming a single cluster.

### STARTS in collinear blocks - A spatial pattern conservation of START genes among ten Oryza species

Occurrence of several genes into a collinear block provides clues on the spatial conservation of the individual genes and their proximal neighborhoods that provides biological significance of gene blocks in the evolutionary sense. Figure 4 depicts these blocks as maps with START genes within one block linked by domain structural classes and collinear gene sets vary from 12-20% across the genome, majority being close to 15% (Supplementary Table S3).

**Figure 4.**
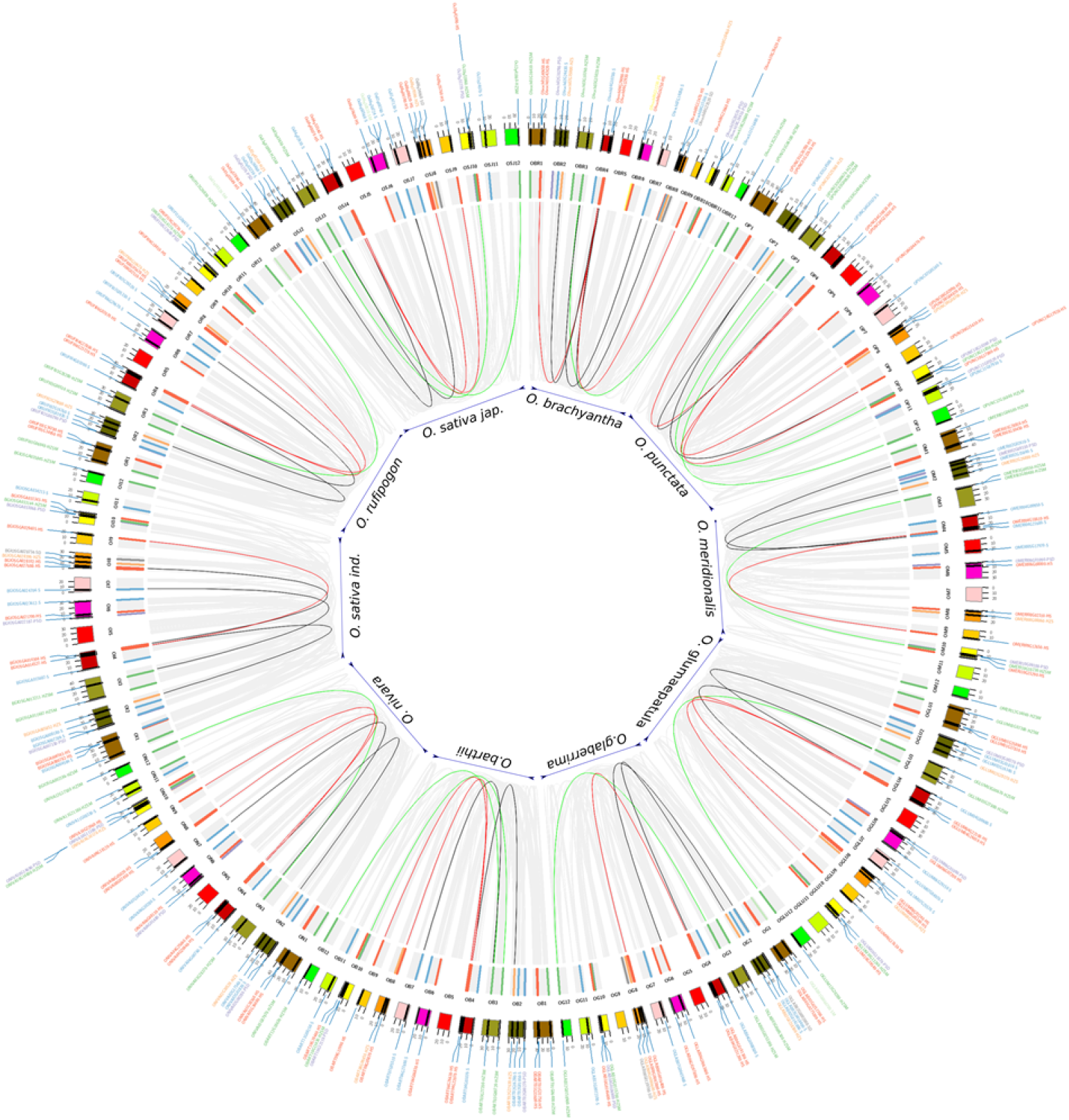
Self-collinear blocks of ten Oryza species. Outer circle: Twelve chromosomes from each of the 10 Oryza genomes that are represented in default colors of circos. Inner circle: START genes are shown as colored bar highlights. Color codes for bar highlights and labels are same as Figure 2&3. Connectors (internal to the bar highlights) are formed between the SART homologues that occur as collinear blocks on different chromosomes of the same genome and follows same color codes of START types except for START homologs that belongs to two different structural classes are linked with a black line. Grey connectors are used for showing non-START homologs within the genomes of ten rices.

Six to ten START genes occurs in collinear blocks, with about one third gene number increase in the cultivated varieties *i.e. Ojap_c_* and *Oind_c_* as compared to early evolved rice species *Obra_w_*, *Opun_w_* or *Omer_w_* and similar is the case of total gene numbers between AA genotype rice varieties *Ojap_c_* (recently evolved) and *Omer_w_* (early evolved). Lack of consistency in the number of genes in syntenic blocks hints at possible chromosomal rearrangement of fragments bearing START genes in both wild and cultivated rice species. The patterns of collinear blocks of cultivated varieties *Ojap_c_* appear similar to immediate ancestors *Oruf_w_*, unlike *Oind_c_*, each pair being placed next to each other in Figure 4. In contrast, the number of START genes in cultivated varieties is reminiscent of their wilder early evolved relatives, for instance, cultivated variety *Ojap_c_* has ten START genes in collinear blocks, similar to its indirect/wilder ancestors *Obar_w_* and *Opun_w_*. The African domesticated varieties *Ogla_c,_* has seven START in its collinear blocks, equivalent to its wilder relative *Omer_w_*. Supplementary Table S2 provides species-wise total number of collinear blocks and START genes that occur in these blocks, while individual circos maps of syntenic collinear blocks are in Supplementary Figure S2 A-J.

### Identification of different modes of START gene duplication

In plants, whole genome duplication leading to polyploids is a frequent event, gene duplication being an important evolutionary phenomenon that helps in the gene dosage, adaptation and speciation; common modes being segmental duplication (SD), dispersed duplication (DD), tandem duplication (TD), and transposed duplication (TsD). Different modes of gene duplication were analyzed for START genes across ten cultivated and wild Oryza species and revealed START genes to exist as duplicated pairs as shown in Table 4. As can be seen in this Table, START genes are rarely present as singletons; and there are two major modes of gene duplication, namely, dispersed and segmental (arising from WGD) across the ten species. Interestingly, dispersed and segmental duplications are similar between pairs of cultivated rice species and their immediate ancestors *(Ojap_c_ and Oind_c_ with Oruf_w_; Oglab_c_ with Obar_w_)* but the proximal and tandem START genes appear to have duplicated after speciation, as the immediate ancestors do not have any. Proximal and tandem duplicate modes among START genes are observed in only two of the early evolved wild species (*Omer_w_ and Opun_w_).* Figure 5 shows a START gene dendrogram with various modes of duplication and paralogous pairs for the main cultivated variety *Ojap_c_*, and it can be seen that of the eight pairs of duplicates, five pairs are segmentally duplicated (between chromosomes 2, 3, 4, 8, 9, 10, and 12), while the three START genes (all on chromosome 6 – see Figure 5) are proximally duplicated. Besides these, two additional STARTS that are found in newly transposed locations as compared to their ancestral gene locations (Figure 5).

**Table 4.**
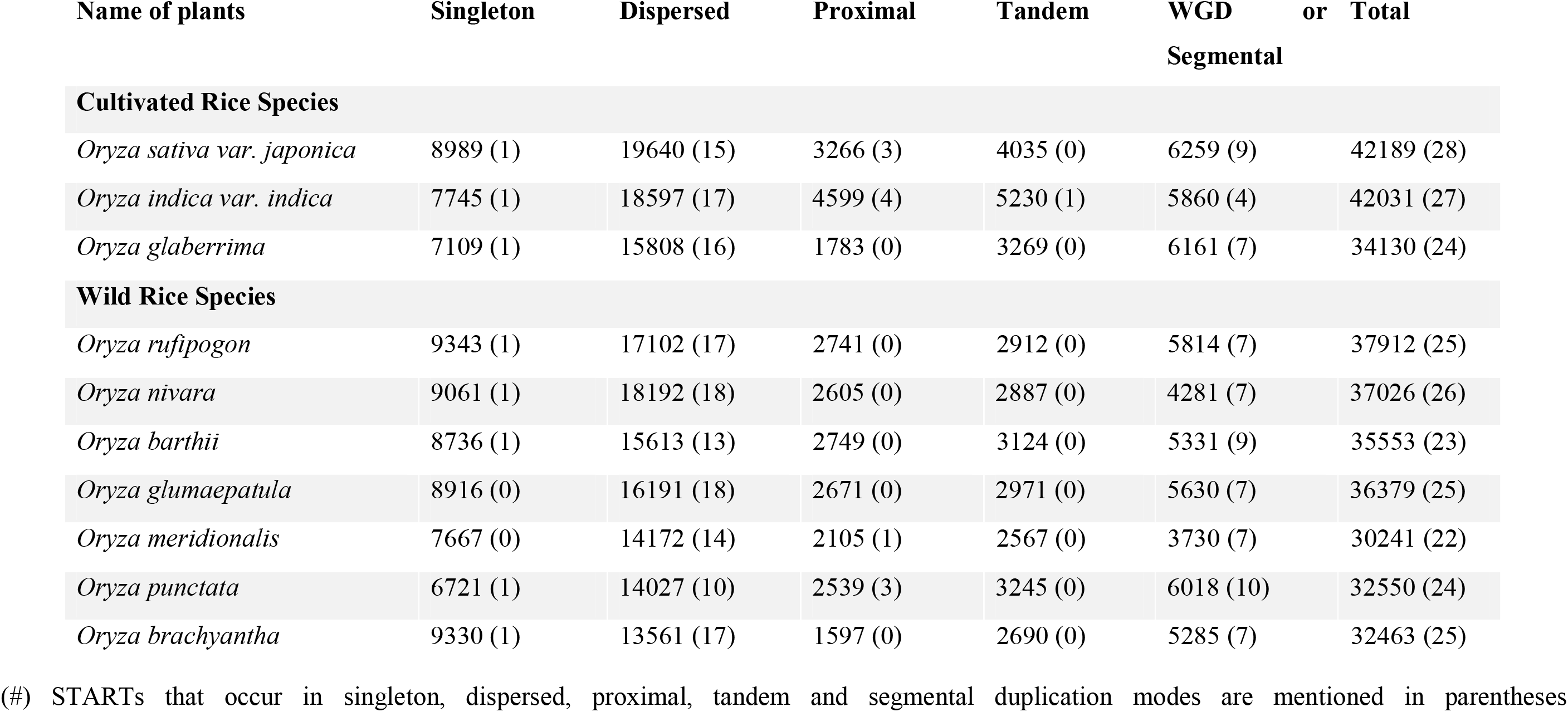
Distribution of different modes of Gene duplication based on whole genome and START genes (in parentheses) amongst ten cultivated and wild Oryza species.

**Figure 5.**
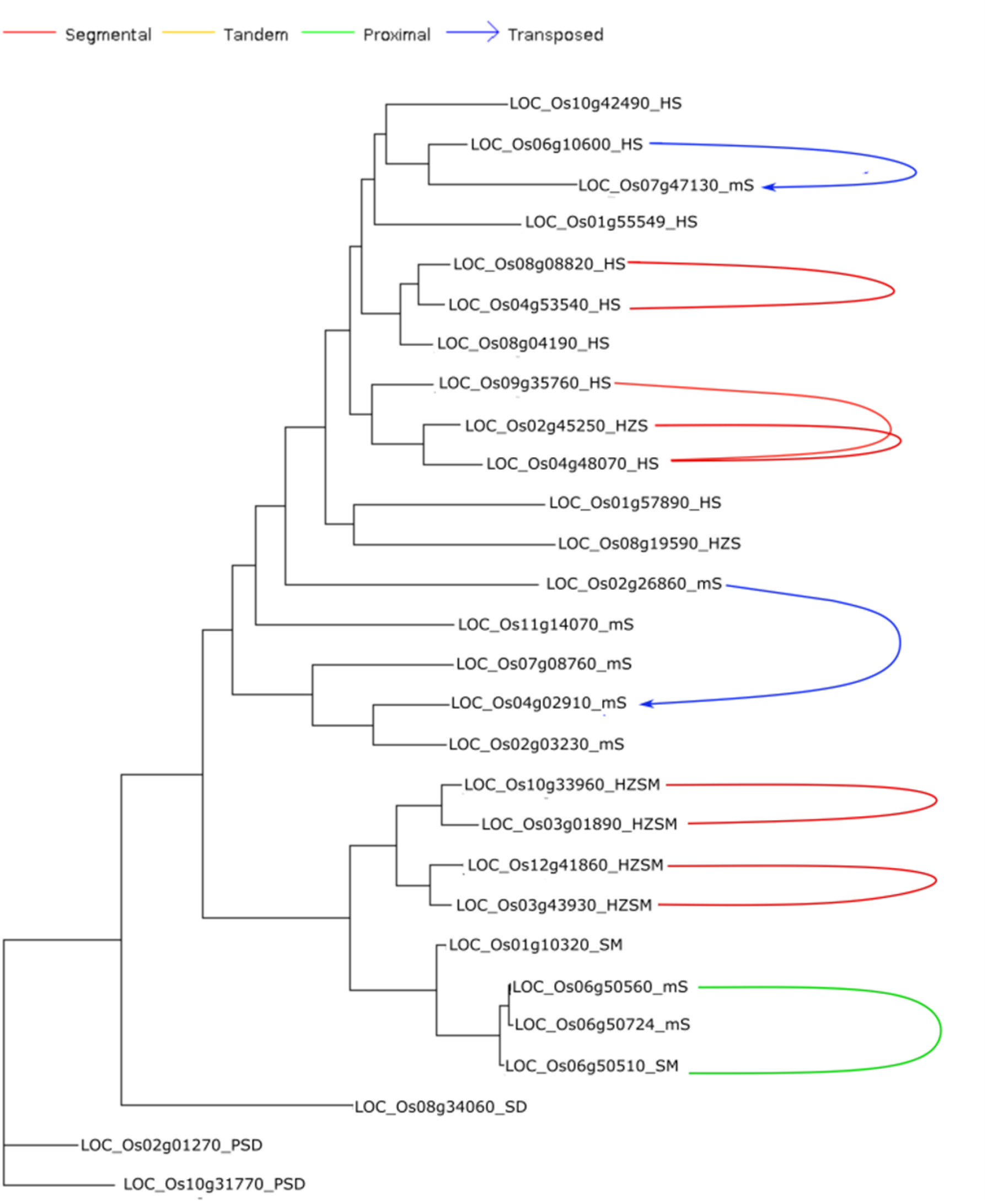
Phylogenetic tree of the *Oryza sativa var. japonica* START genes family annotated with different modes of gene duplication

### Nucleotide Substitution Rates and Ka/ Ks Ratios

Ka/Ks ratios represent selection pressure on genes, with values of >1, <1, or 1 signifying positive, negative or neutral selection, respectively [55]. These calculations for START genes of all nine Oryza genomes with respect to recently evolved cultivated variety *Ojap_c_* as described in methods are shown in Figure 6 panels A-E, and Supplementary Figure S3A-C. With few exceptions, most of the START gene pairs have Ka/Ks values below one, suggesting their being under negative selection. The unique domain categorical group of ‘SM’ in *Ojap_c_* and its orthologous pairs in *Oind_c_*, *Ogla_c_*, and *Oniv_w_* showed a very high positive selection suggesting their being under positive selection. Apart from this there are few other cases, which also showed Ka/Ks values significantly more than 1, and close to 1, which signifies that these START genes are also undergoing through positive selection. The analysis further confirmed a high rate of synonymous and non-synonymous substitutions for both the PSD type orthologs and single HS homologue (present on chromosome 4).

**Figure 6.**
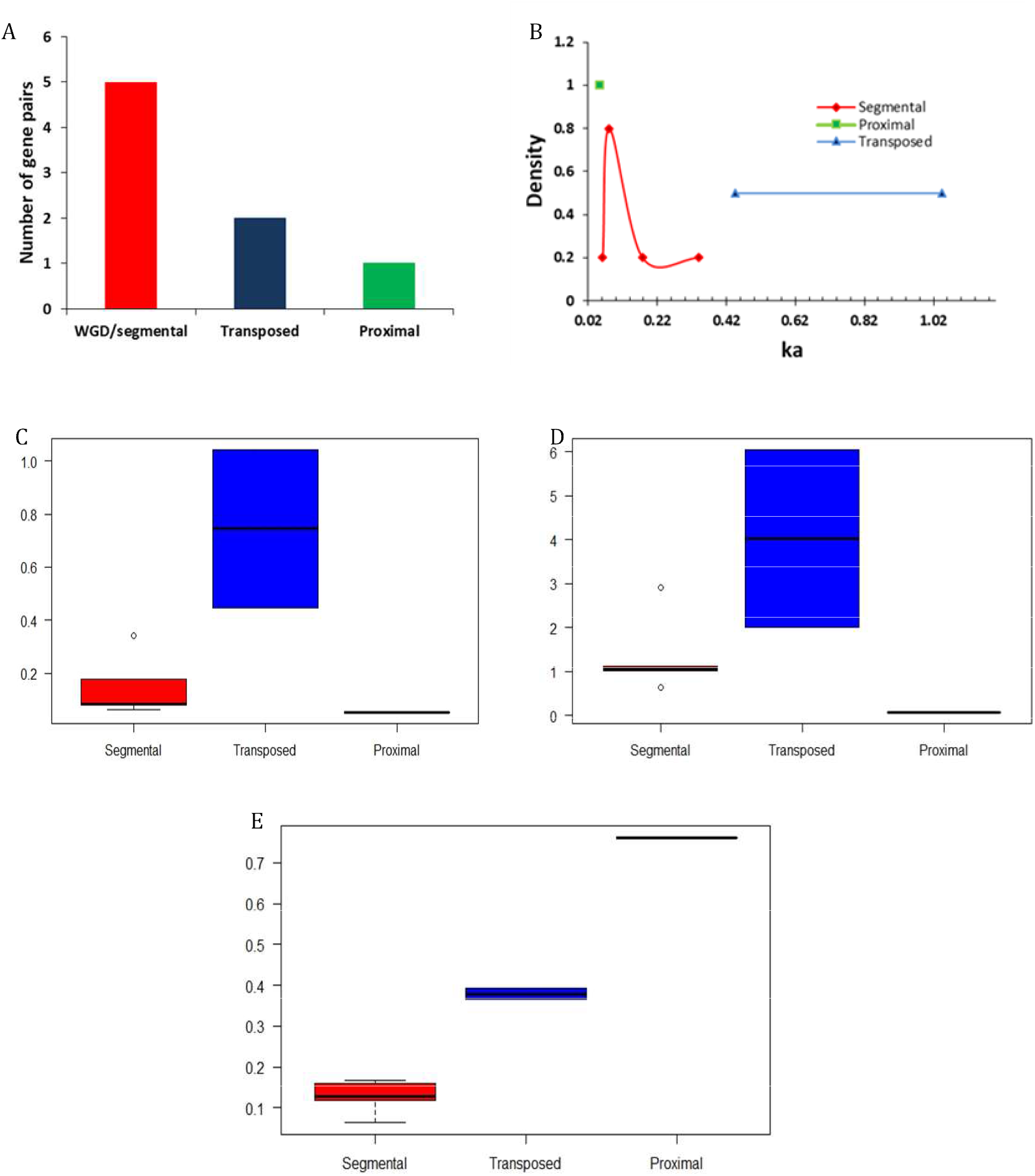
Evolutionary patterns of duplicated START gene pairs by different modes in *Oryza sativa var. japonica.* (A) Gene pair distribution among different modes of duplication. (B & C). Ka values of duplicated gene pair (D). Ks value of duplicated gene pairs (E). Ka/Ks values of duplicate genes pairs

The eight paralogous START gene pairs of *Ojap_c_* that were noticed in segmental, transposed and proximal modes of duplications (Figure 5) were further evaluated for the Ka, Ks, and Ka/Ks. As shown in Figure 6, all gene pairs have negative Ka/Ks ratios suggesting low or moderate flexibility for mutational changes. One proximal duplicate pair (LOC_Os06g50560_mS-LOC_Os06g50510_SM) at 0.762 suggests flexibility in mutational rate of the second copy gene, whereas for the five segmental START gene duplicates, Ka/Ks varied between 0.166 to 0.063, indicating stringent regulation and highlighting their functional importance (LOC_Os10g33960_HZSM-LOC_Os03g01890_HZSM, LOC_Os12g41860_HZSM-LOC_Os03g43930_HZSM, LOC_Os08g08820_HS-LOC_Os04g53540_HS, LOC_Os04g48070_HS-LOC_Os02g45250_HZS, and LOC_Os04g48070_HS-LOC_Os09g35760_HS) (Supplementary Table S4). As presented in the next section, we also observed similar expression levels among these pairs across anatomical parts but with slight variation between different stages of development.

Overall, Ka/Ks analysis of duplicated START gene pairs in *Ojap_c_* suggest that the expanded START gene family is still evolving towards stabilization of function and expanding into new roles or sub-functionalization. The Ka/Ks results also suggest that proximally duplicated STARTs have 99% of identity amongst themselves and have not undergone changes, compared to other modes of duplications, which might be due to the recent incidence of this mode of duplication. This is also supported by the phenomenon of the overall number of accumulated mutations over the evolutionary history of an organism [56]. Contrastingly, the transposed duplicates underwent intermediate negative selection pressure and the segmental duplicates underwent strong negative selection pressures. Similarly, the synonymous substitution rates for transposed duplicates were higher when compared to segmental START duplicates (Supplementary Table S4 – The transposed pair LOC_Os02g26860_mS-LOC_Os04g02910_mS showed 3 fold higher Ka and Ks values supporting the phenomenon of evolutionary freeness for development of sub-functionalization or neo-functionalization when compared to segmental duplicate gene pair *i.e.* LOC_Os04g48070_HS-LOC_Os09g35760_HS, which showed similar Ka and Ks values with other transposed pair LOC_Os06g10600_HS-LOC_Os07g47130_mS showing slight stringency in mutational frequency of these genes.

The transposed pairs (Os02g26860_mS-LOC_Os04g02910_mS and LOC_Os06g10600_HS-LOC_Os07g47130_mS; above 0.365 Ka/Ks ratio) in addition to the proximal duplicated pairs (LOC_Os06g50560_mS-LOC_Os06g50510_SM; 0.762 ka/ks ratio) had the highest mean Ka/Ks ratio indicating that they have experienced weaker purifying selection. The segmental gene pair (LOC_Os10g33960_HZSM-LOC_Os03g01890_HZSM) had the lowest mean ka/ks ratio (0.063) suggesting strong purifying selection and the other four segmental pairs (LOC_Os12g41860_HZSM-LOC_Os03g43930_HZSM, LOC_Os08g08820_HS-LOC_Os04g53540_HS, LOC_Os04g48070_HS-LOC_Os02g45250_HZS, and LOC_Os04g48070_HS-LOC_Os09g35760_HS) with intermediary mean Ka/Ks ratio above 1, indicating that they had experienced intermediate to stronger purifying selection. Thus, START genes appear to be under purifying selection pressure, further highlighting their functional importance and roles for expansion of START genes among wild and cultivated rices, and we explore this further through gene expression analyses.

### Transcriptome analysis of START encoding genes

The function of many START genes especially Homeodomain associated START genes have been extensively studied in plants. The class III HD-ZIP family and classes IV HD-ZIP family have a well established role in Arabidopsis and involved in various stages of development and gene regulation [8]. In order to explore the potential functions of the 28 START genes found in *O. sativa var. japonica,* the tissue and developmental stage specific expression pattern were investigated in non-stressed condition as described in methods. Figure 7 depicts the expression heat maps of START genes, four HZSM, two HD bZIP STARTs and two START MEKHLA genes in *O. sativa var. japonica* showed significant expression throughout the developmental stages and anatomical parts. As can be seen in Figure 7A, five out of nine HD-START genes express constitutively through the various developmental stages, but almost all of these nine genes show differential expression across various anatomical parts (Figure 7B), suggesting tissue specific roles for this large amplified sub-group. The eight minimal START genes (LOC_Os02g03230_mS, LOC_Os02g26860_mS, LOC_Os04g02910_mS, LOC_Os06g50560_mS, LOC_Os06g50724_mS, LOC_Os07g08760_mS, LOC_Os07g47130_mS, and LOC_Os11g14070_mS) of *Ojap_c_* show a wide variation in expression patterns across all stages and tissues, and it is possible to assign them to tissue specific roles, with only one (LOC_Os07g08760_mS) showing high expression across all anatomical parts. START genes were grouped via hierarchical clustering of expression profiles and this is depicted in Figure 7 panels C and D. Comparison of this data with duplication analyses in the earlier section reveals that most of the duplicated START gene pairs show similar expression pattern across both developmental stages and anatomical parts, with the exception of proximal duplicated START genes (detailed analysis in Supplementary Table S4.). Expression patterns among five segmental gene pairs varied with three pairs in one cluster and two in other clusters, despite showed significant expression throughout all the developmental stages. The five segmental pairs constitute two pairs of HS genes (4 genes) and this cluster together in expression as described in the next section. Taken together (Fig 5 &7) duplicated START genes in segmental and transposed modes showed a unified pattern in gene expression amongst duplicated gene pairs across all the developmental stages as well as in all anatomical parts, which signifies the functional importance of STARTs and also the necessity of retaining both the copies of the gene pairs. Contrastingly, the proximal duplicate gene pair showed an uneven expression pattern between the gene pairs, which indicates a possible adaptation towards sub-functionalization or neo-functionalization of the duplicated gene, as was observed for the Ka/Ks selection pressures.

**Fig. 7.**
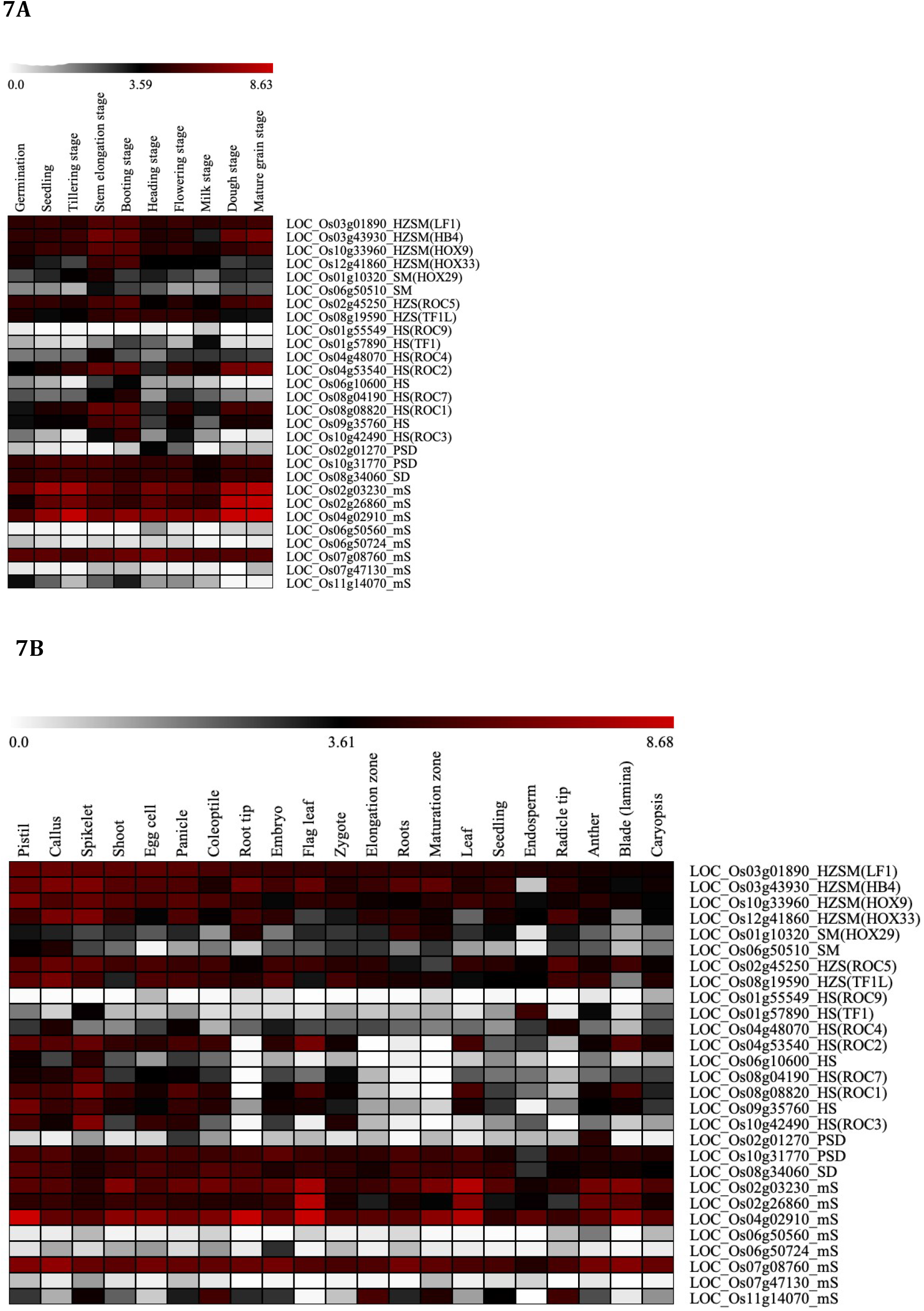
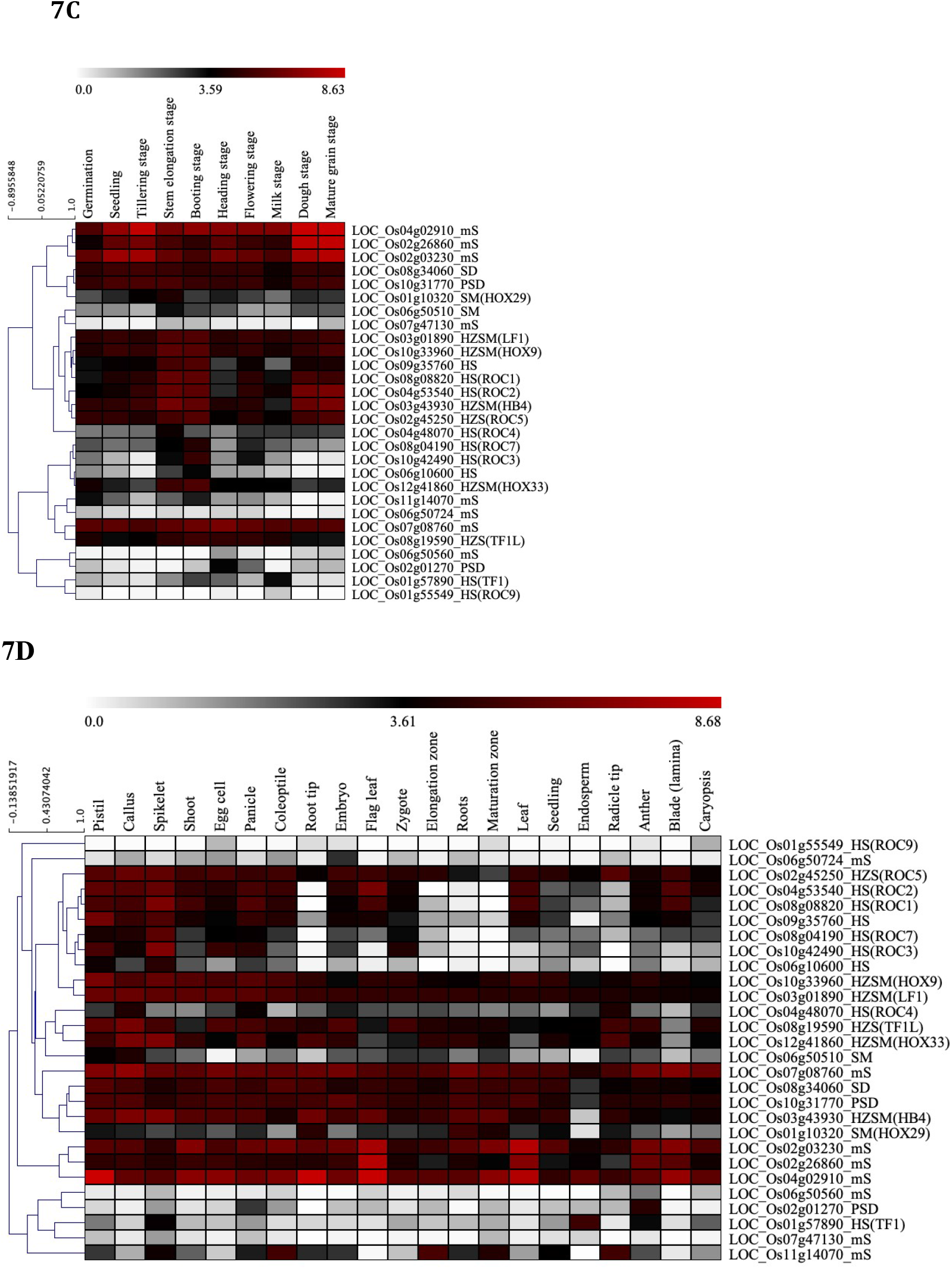
*In-silico* expression study of 28 START genes in *Oryza sativa var. japonica.* A) Conditional gene expression pattern of START genes at Different developmental Stages B) Conditional gene expression pattern of START genes in various anatomical parts C) Hierarchical clustering of START genes for developmental stages D) Hierarchical clustering of START genes for anatomical parts Common name in brackets, wherever available.

## Discussion

Genome duplication events play a significant role in environmental adaptation and speciation of organisms. Studying the function of post duplication genomes has shed light on how genomes evolve and their functional diversification. Loss of alternative copies of the duplicated locus led to reproductive isolation between mating populations. The major changes observed at the protein level in organisms, in the post-genome duplication events, are copy gain or loss and domain alterations, i.e. gain or loss of domains and in some cases domain rearrangements that occur in the new copy at the gene level which decides the adaptability of organisms to the challenging environments. This phenomenon has been reported among all the kingdoms of life due to dosage of the extra gene product, and its spatio-temporal expression changes [57–60].

Over a decade ago, it was first revealed that START genes are amplified in plants compared to animals and this led to an era of interesting work in this area to understand why these domains are amplified, and what roles they may be performing in plants [6]. Presence of this domain among evolutionary distant organisms such as plants, animals, a well as some species of bacteria and protists exemplifies this domain’s evolutionary significance throughout different domains of life [6,61]. Humans and mouse are having 15 START protein encoding genes [4]. While a comparative study of START domains in *Arabidopsis thaliana* shows that it has 35 START genes, whereas rice is having 29 START gene orthologs, cumulatively in Indica (having 27 genes) and Japonica (Having 22 genes) subspecies [6]. The increase in number of START genes (28) in *Ojap_c_* compare to previous studies is mainly because of a newer version of genome assemblies (MSU Release 7.0) is used in this study. Many primarily studies were carried out regarding the plant START proteins mainly Homeodomain (HD) associated START proteins in plants [6,18,62,63] however this study, first to our knowledge, reports the comprehensive comparison of START genes amongst the inter and intra-genuses of wild and cultivated species of rices. Our genome-wide comparative studies shows that START gene family size slightly varies among the cultivated and wild rice species. START proteins are found as single domain bearing minimal START proteins, as well as multi domain proteins containing one or more domains of known or unknown functions such as HDs, MEKHLA, PH, and DUF1336 and this classifies the START proteins into many structural classes. Moreover, orthologous studies among ten rice species emphasizes that, the major structural classes of START genes are preserved across the wild and cultivated rice species, although with a minor differences in their numbers. Additionally, this study advocates that, the different structural classes of START proteins may have a specified functional role in plants.

Gene duplication events have led to the expansion of the START gene family in Oryza genome, as duplication studies show that almost 95% of START genes are duplicated, mainly through Dispersed and Segmental modes of duplication. Additionally, the regulation of Ka/Ks ratio in the intermediate level for segmentally duplicated START gene pairs highlights their significance in functional roles. The lower stringency in the Ka/Ks ratio for transposed and proximal START gene pairs indicates the incomplete sub-functionalization or neo functionalization of these START genes in Oryza sativa japonica genome. Similarly, Qiao et al. 2019 have reported that above 98% of the genes in the oryza sativa genome underwent purifying/negative selection[64]. They have also showed that below 2% of tandem, proximal and dispersed gene pairs underwent positive selection, while 0.5% of the segmental and transposed duplicate gene pairs in Oryza sativa genome underwent positive selection. Additionally, their result shows that higher proportion of tandem and proximal duplicated gene pairs showed gene conversion in rice and Arabidopsis genomes than the duplicated gene pairs through other modes of duplication mechanism. Our results of Ka/ks ratio estimates for both transposed START duplicate pairs showed positive selection which may fall into the exceptional category of transposed duplicates (below 2%) that Qiao et al. 2019 have mentioned, that showed positive selection, while the *Ojap_c_* proximal START duplicate results completely agree with their results in *Orya sativa var. japonica* [64].

There are very few reports available on the protein domain level changes post-genome duplications. Finet et al. 2013 showed that gene truncations, separating the DNA-binding fragment from the response fragment, in one of the copies of Auxin response factor (ARF) gene duplicates that resulted in the structural and functional diversity of the ARF proteins[65]. They have also proposed that at least two independent events of truncations that led the formation of two distinct ARF3 subclades in monocots, which lacks either both domains III, and IV or only domain IV. These earlier reports supports our data where formation of novel START classes such as PS and SM is only possible due to domain gain/loss mechanism. The evolutionary significance of these novel START protein structural classes was obtained from their significant level of expression during various anatomical and developmental stages..

Our current results also suggests the possibility of two independent truncation events that may have led the formation of SM subclades of START proteins either from HZSM clade or mS clade. There may be a possible domain gain/loss in the PSD structural class leading to PS subclade or vice versa. Kersting et al. (2012) report on the domain loss in monocots with their studies on plant domain analysis (average domain loss rate of 7.4/Myr) in 29 plant genomes and 3 algal genomes strengthen the idea of possible domain loss mechanism in certain classes of START proteins to form novel structural classes [66]. Additionally, their data have also showed that there were higher domain gain/loss events in *Oryza sativa* genome than *Brachypodium distachyon*. These all earlier studies have supported the domain level changes gain, loss or rearrangements are involved in the adaptability of the plants the environmental changes. Overall, our Phylogenetic, gene duplication, synteny, nucleotide substitution rates and transcriptome analysis of START genes in rice genome, highlights the different gene amplification mechanism, their functional significance and evolutionary changes that may have led to the formation of new structural classes.

## Conclusions

START domains are abundant in plants and play a crucial role in plant physiology and development. In this paper we have identified START family proteins in ten Oryza genomes and classified these into structural classes based on additional START-domain associated functional domains, further phylogenetic analysis confirmed significant evolutionary divergence among these structural classes. We overlay the analysis onto the wild and cultivated rice varieties and bring our features relating to evolution and ancestry of these domains within the ten species investigated, in order to understand the causes and consequences of START gene family expansion. Patterns from analysis of selection pressures were superimposed on gene expression profiles for the most widely used rice variety across the globe, namely *Oryza sativa var. japonica*, further confirming functional importance of the large expansion of this important gene family in plant development and tissue specific roles. We hope this work on START gene family in Oryza species will pave the way for exploring the functional mechanism and substrate preference of plants START domains.

## Patents

Not Applicable

## Author Contributions

The work was conceived by GY, research work was performed by SKM & RKP. All three authors performed data analysis, wrote the manuscript and approved for final publication.

## Funding

SKM receive fellowship from the Department of Biotechnology (DBT) Government of India and National Institute of Plant Genome Research (NIPGR) during his Ph.D (Fellow no. DBT/2014/NIPGR/265). RKP is funded by the Department of Biotechnology, Government of India [Grant ID BT/PR22334/BID/7/786/2016].

## Acknowledgments

Authors acknowledge the support of National Institute of Plant Genome Research (NIPGR), New Delhi for infrastructure and DBT-eLibrary Consortium (DeLCON) for providing access to e-resources.

## Conflicts of Interest

None

## Abbreviations

START: StAR-related lipid-transfer
StAR: Steroidogenic acute regulatory protein
HD: Homeodomain
TF: Transcription Factor
START proteins: START domains containing full-length proteins
START genes: START genes that codes for full length START proteins
PH: Pleckstrin homology
DUF1336: Domain of unknown function 1336
HZSM: HD bZIP START MEKHLA
SM: START MEKHLA
HZS: HD bZIP START
HS: HD START
PSD: PH START DUF1336
PS: PH START
SD: START DUF1336
mS: minimal START proteins

## Supplementary Materials

Figure S1A-J: Gene structure analysis of STARTs genes among ten rice species *(A) Oryza sativa var. japonica, (B) Oryza sativa var. Indica, (C) Oryza glaberrima, (D) Oryza rufipogon, (E) Oryza nivara, (F) Oryza barthii, (G) Oryza glumaepatula, (H) Oryza meridionalis, (I) Oryza punctata and (J) Oryza brachyantha.*

Figure S2A-J: Collinear blocks of the ten rice genomes *(A) Oryza sativa var. japonica, (B) Oryza sativa var. Indica, (C) Oryza glaberrima, (D) Oryza rufipogon, (E) Oryza nivara, (F) 912 Oryza barthii, (G) Oryza glumaepatula, (H) Oryza meridionalis, (I) Oryza punctata and (J) Oryza brachyantha.*

Figure S3 (A) Ka, (B) Ks and (C) Ka/Ks values, for START homologues of different Oryza species with respect to *Oryza sativa var. japonica.*

Table S1: The locus ids of START genes along with sequence analysis Information and phylogenetic code.

Table S2: The Detailed gene structure pattern of 10 Oryza genome, 5’UTR, 3’UTR, exon and Intron length.

Table S3. Collinear genes number across ten Oryza genome.

Table S4: Ka, Ks, and Ka/Ks analysis among the gene pairs that follow different modes of duplication in *Oryza sativa var. japonica* genome.

## Notes

### Competing Interest Statement

The authors have declared no competing interest.

