## Supplementary Figure S1A-J, S2A-J, S3A-C for "Comparative genomics of StAR-related lipid transfer (START) domains across wild and cultivated rice"

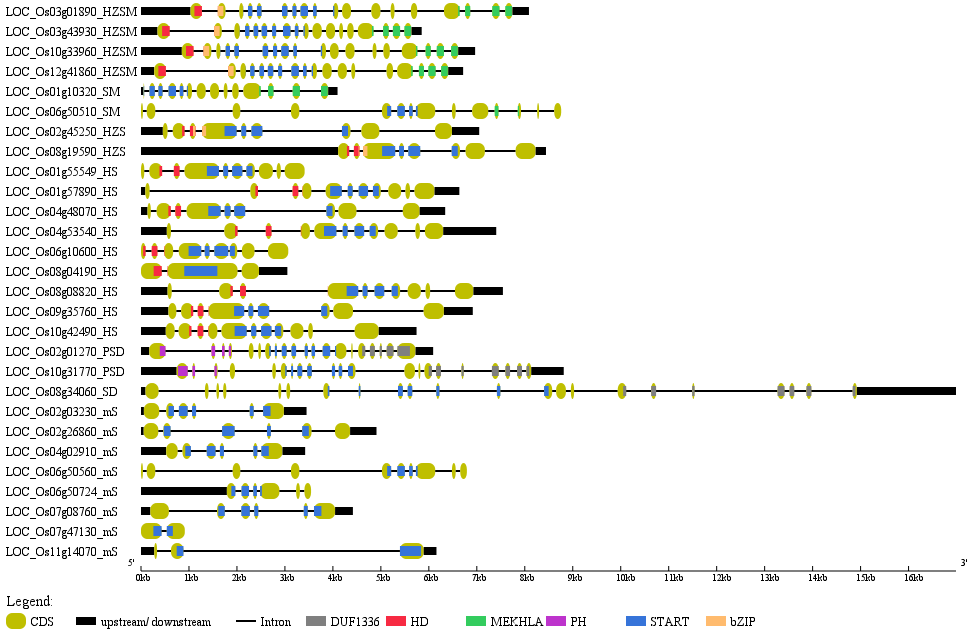

**Supplementary Figure S1A.** Gene structure analysis of the twenty-eight STARTs in *Oryza sativa* var. japonica: exon-intron patterns

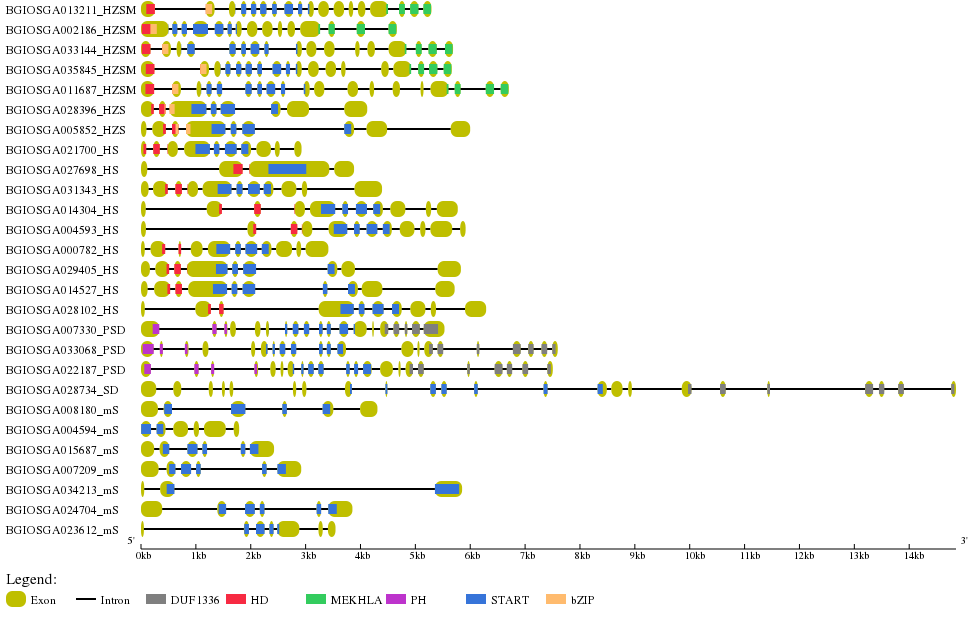
**Supplementary Figure S1B.** Gene structure analysis of the twenty-seven STARTs in *Oryza sativa var. indica*: exon-intron patterns
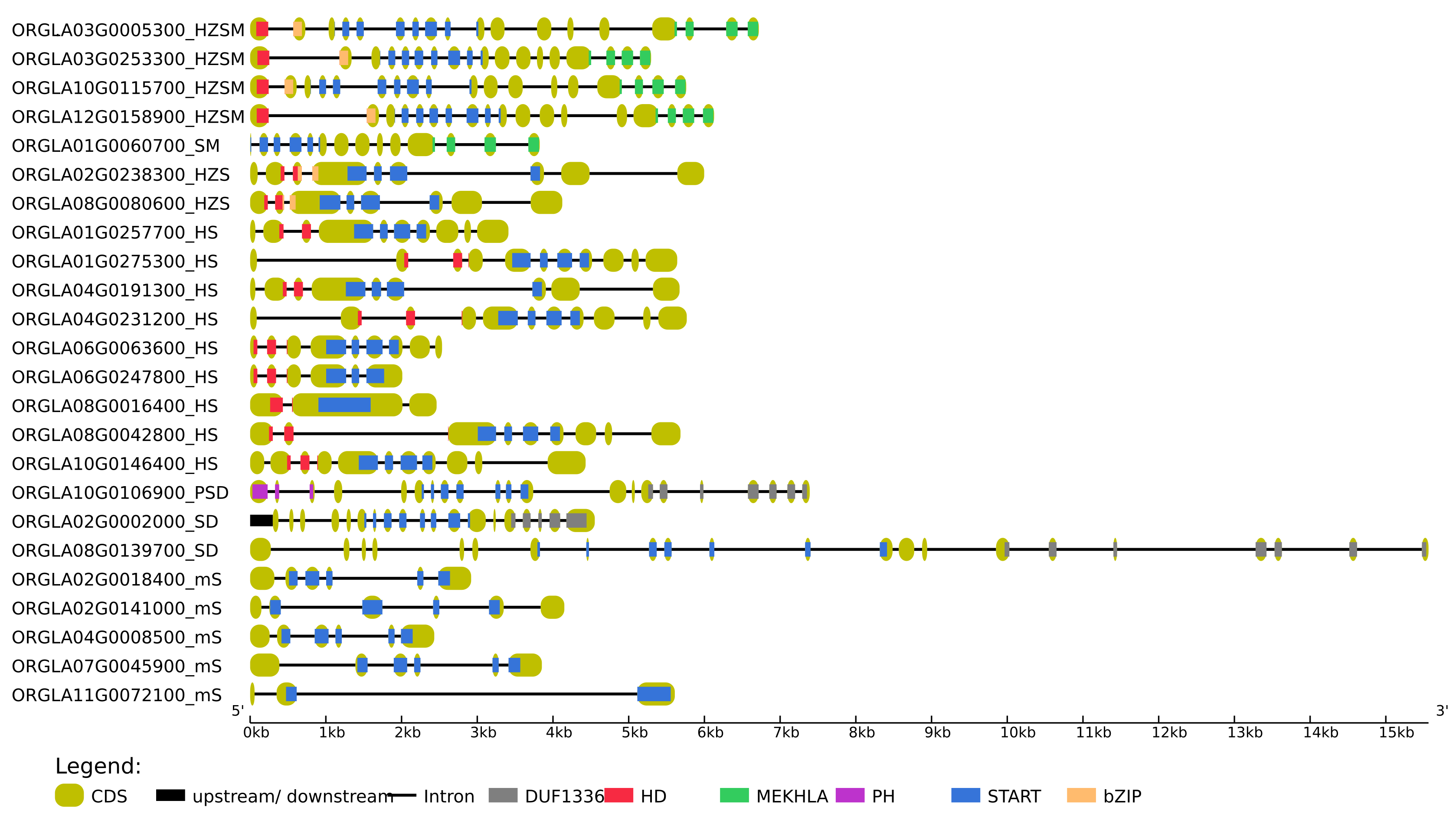
**Supplementary Figure S1C.** Gene structure analysis of the twenty-four STARTs in *Oryza glaberrima*: exon-intron patterns

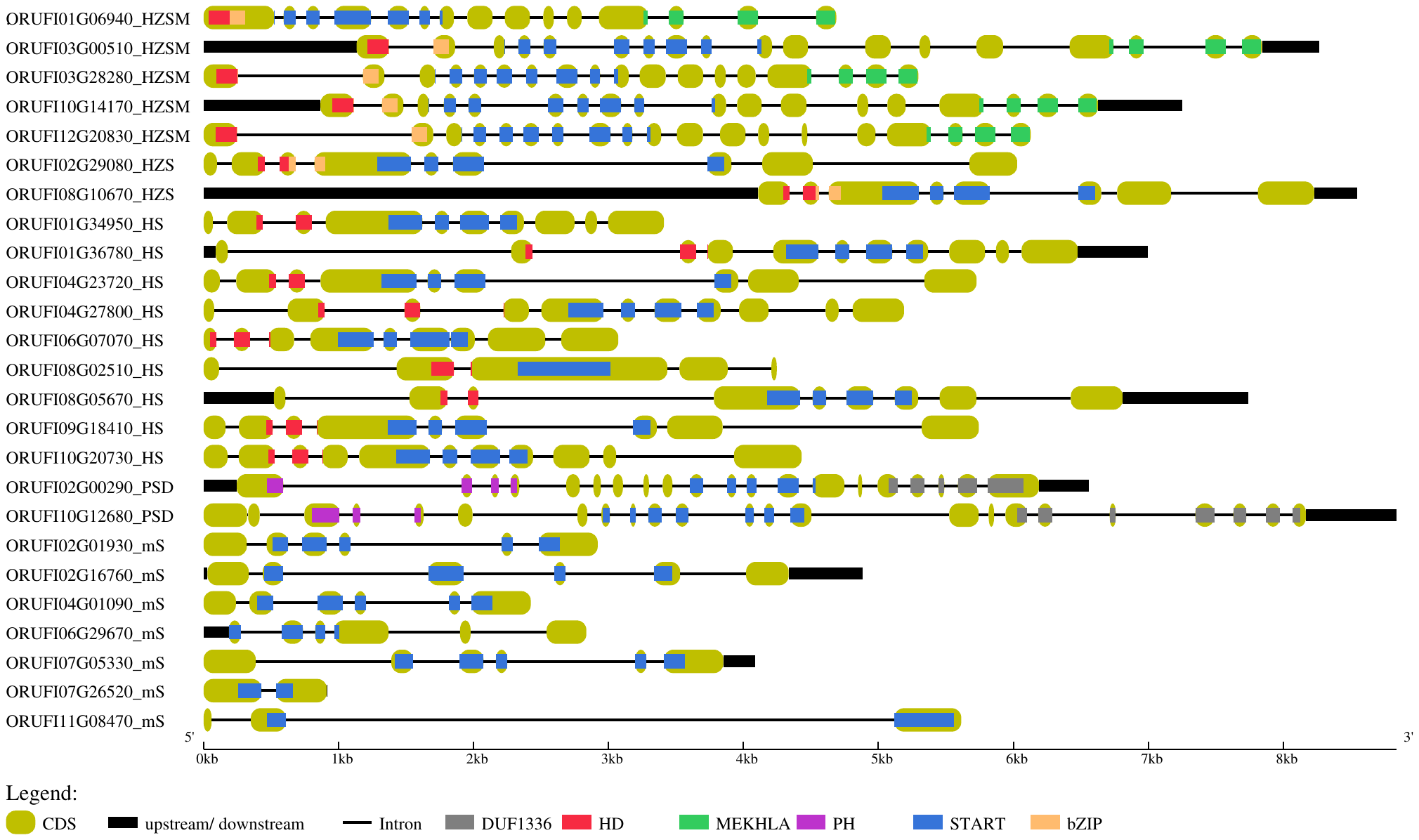

**Supplementary Figure S1D.** Gene structure analysis of the twenty-five STARTs in *Oryza rufipogon*: exon-intron patterns

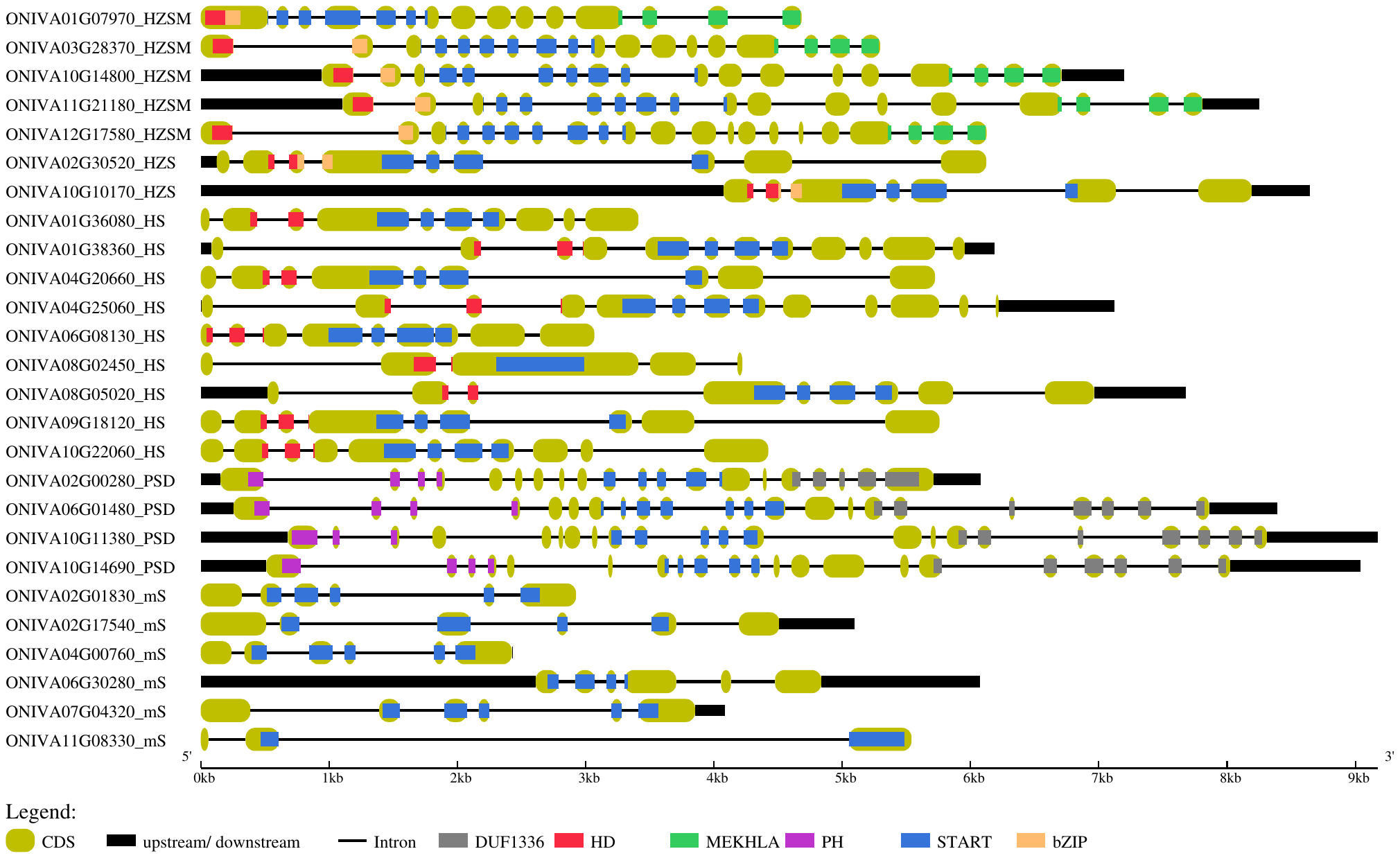

**Supplementary Figure S1E.** Gene structure analysis of the twenty-six STARTs in *Oryza nivara*: exon-intron patterns

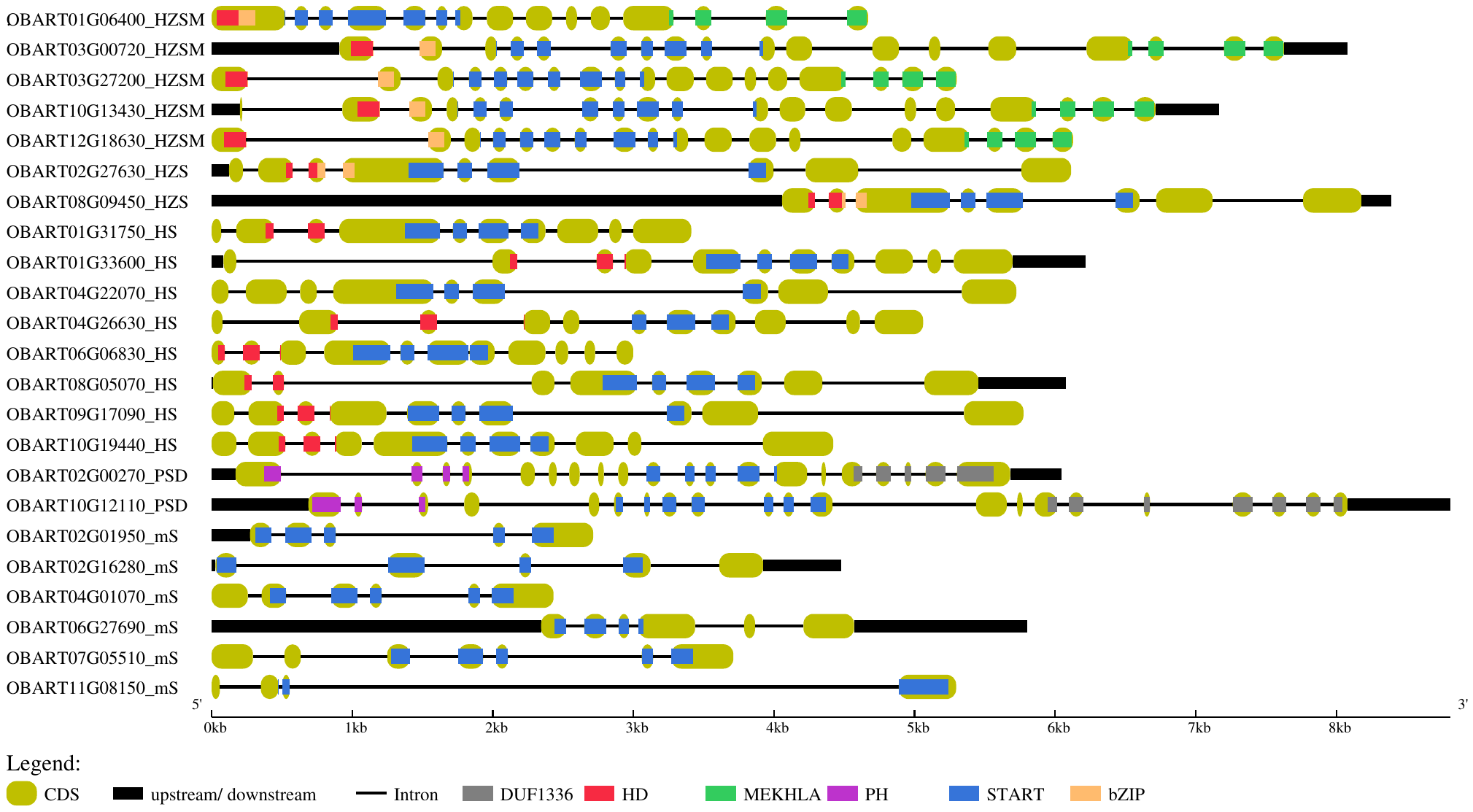

**Supplementary Figure S F.** Gene structure analysis of the twenty-three STARTs in *Oryza barthii*: exon-intron patterns

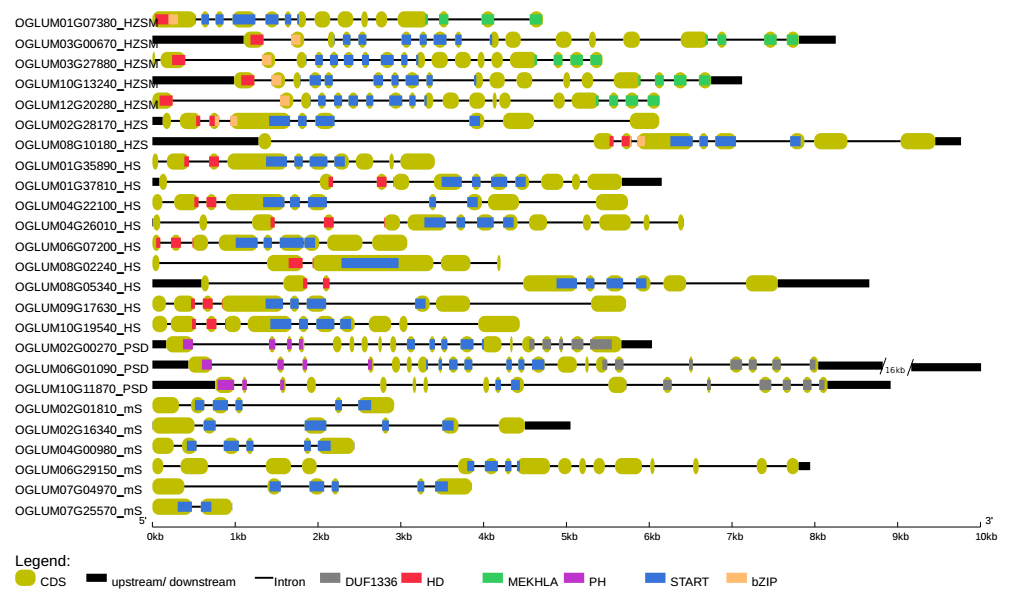

**Supplementary Figure S1G.** Gene structure analysis of the twenty-five STARTs in *Oryza glumipatula*: exon-intron patterns

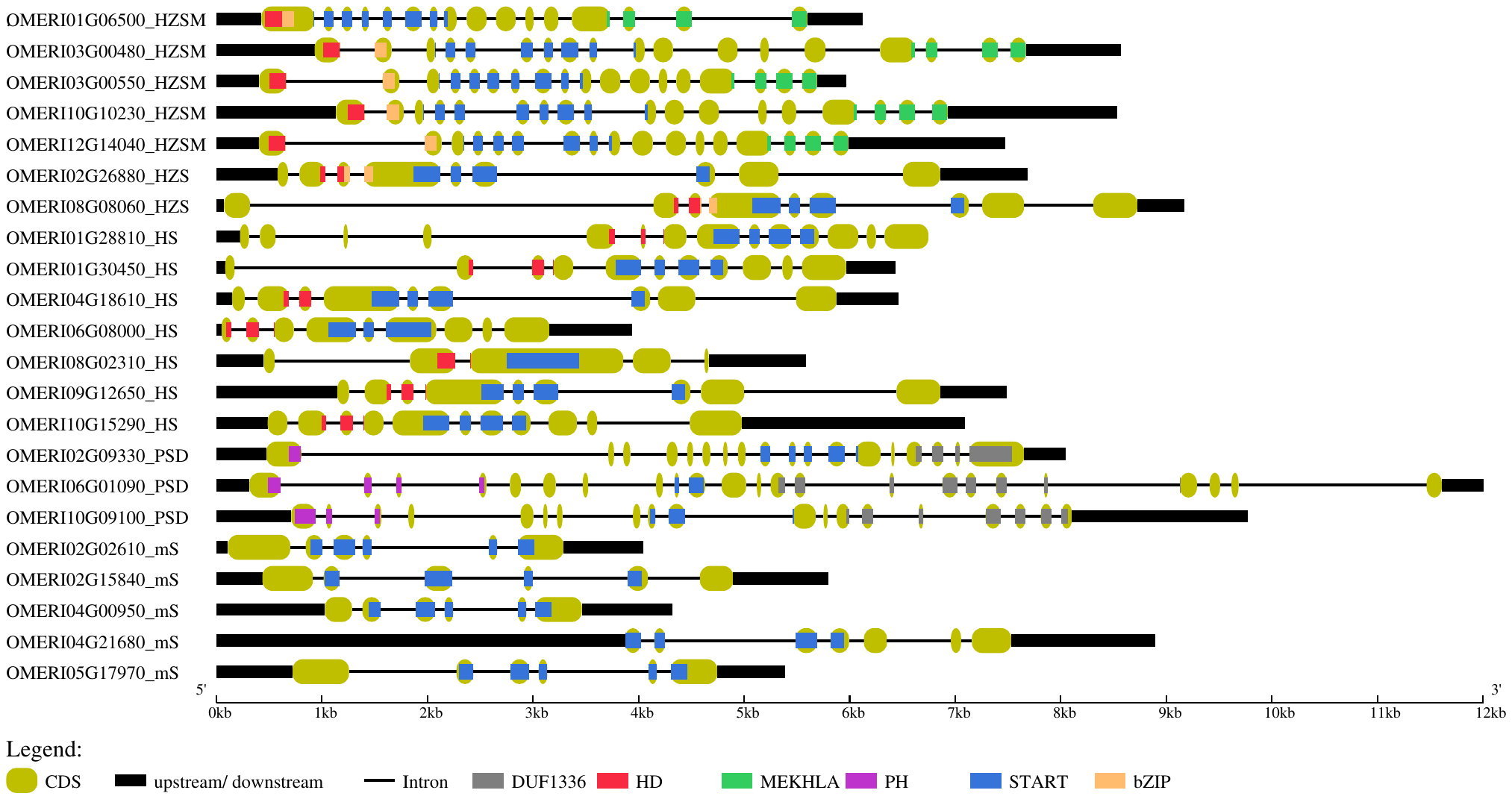

**Supplementary Figure S1H.** Gene structure analysis of the twenty-two STARTs in *Oryza meridionalis*: exon-intron patterns

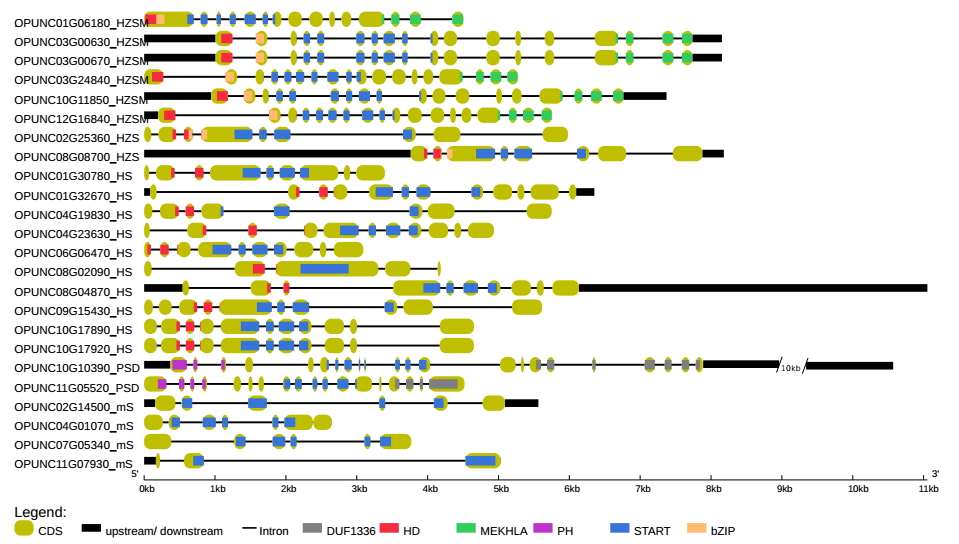

**Supplementary Figure S1I.** Gene structure analysis of the twenty-four STARTs in *Oryza punctata*: exon-intron patterns

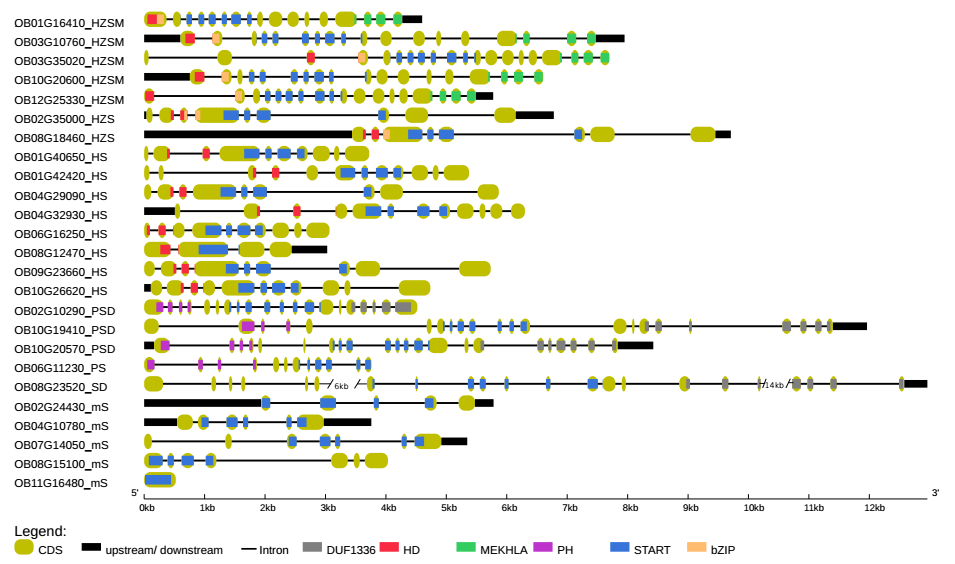
**Supplementary Figure S1J.** Gene structure analysis of the twenty-five STARTs in *Oryza brachyantha*: exon-intron patterns

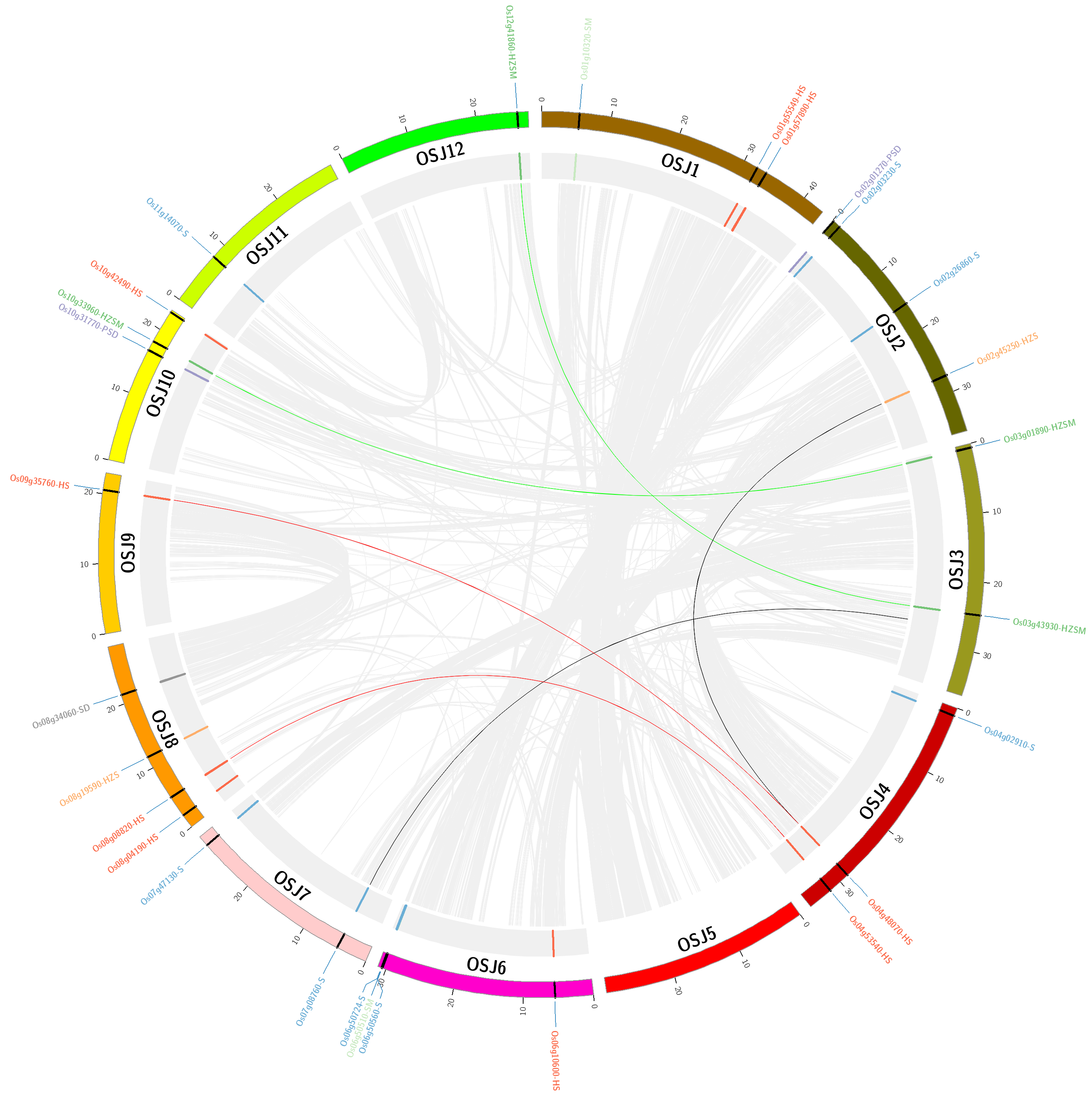

**Supplementary Figure S2A.** Collinear blocks of the *Oryza sativa var. japonica* genome.

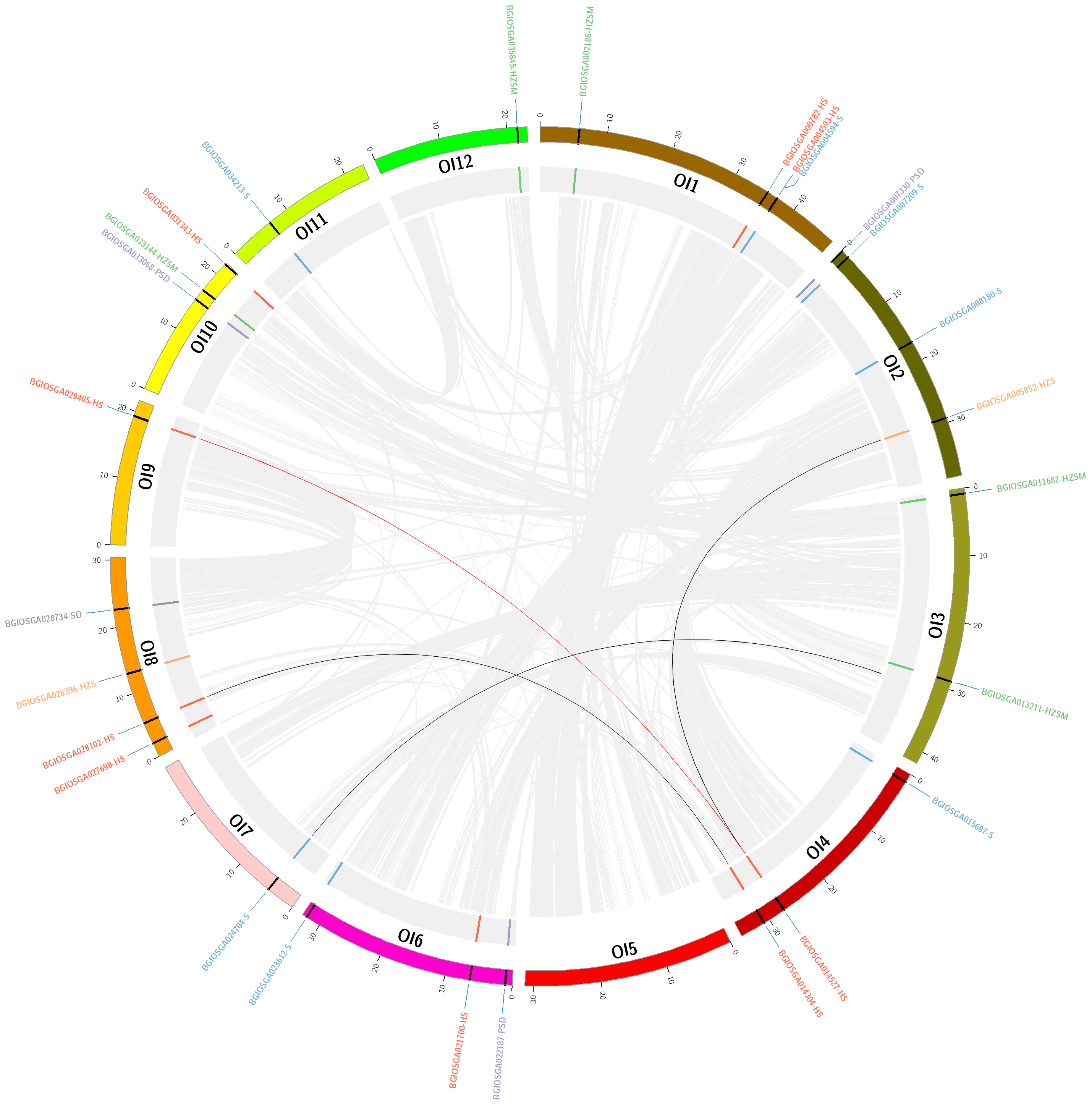

**Supplementary Figure S2B.** Collinear blocks of the *Oryza sativa var. indica* genome.

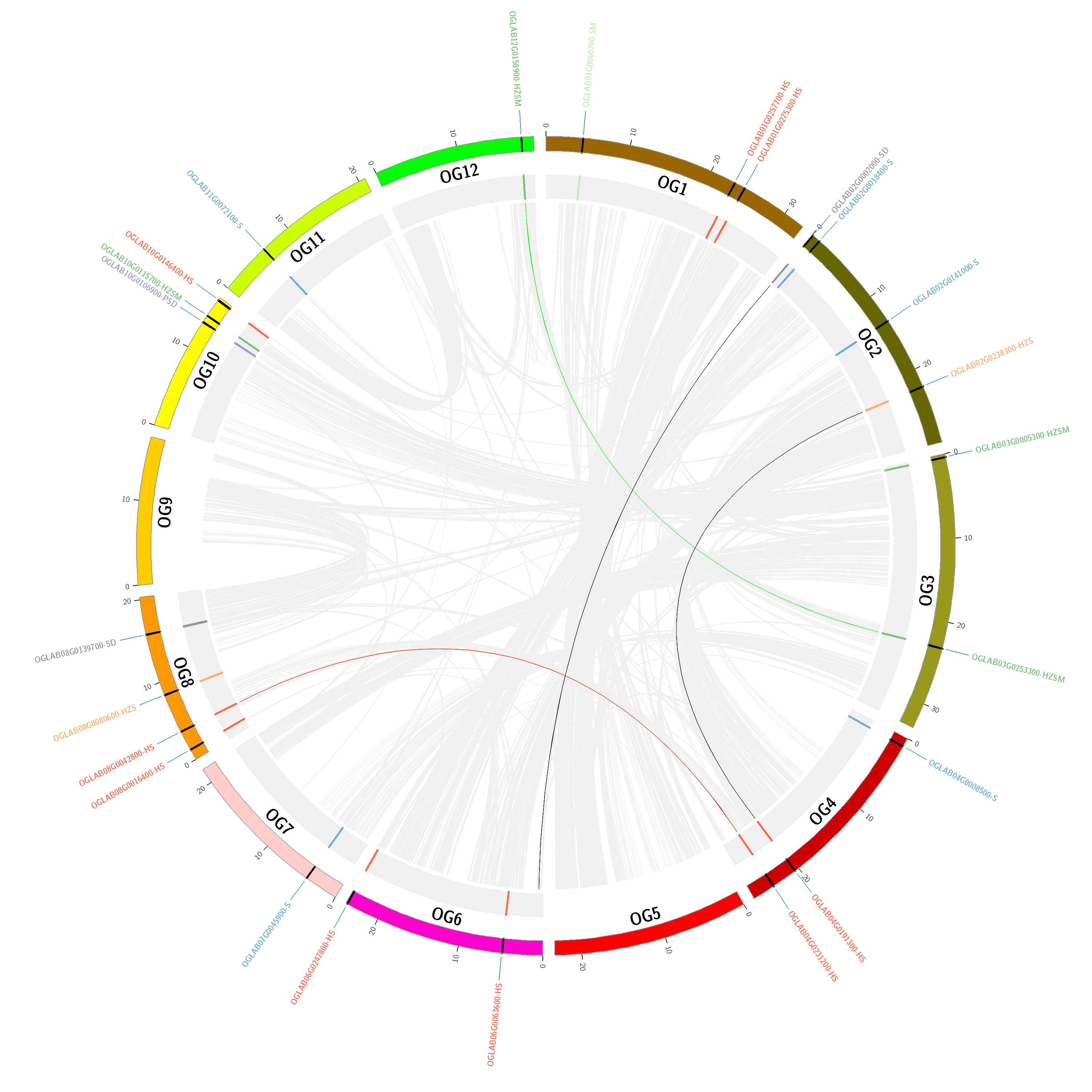

**Supplementary Figure S2C.** Collinear blocks of the *Oryza glaberrima* genome.

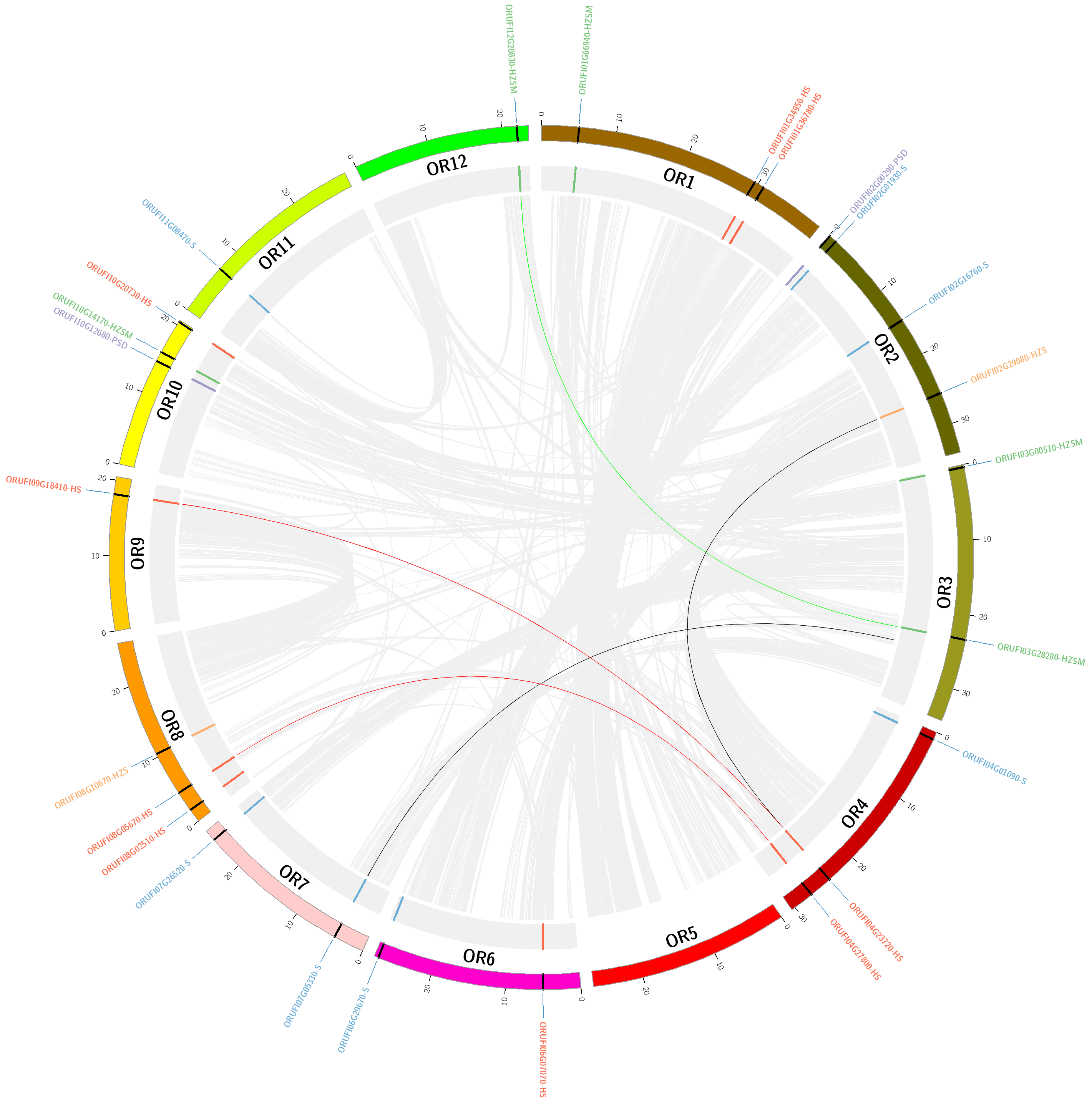

**Supplementary Figure S2D.** Collinear blocks of the *Oryza rufipogon* genome.

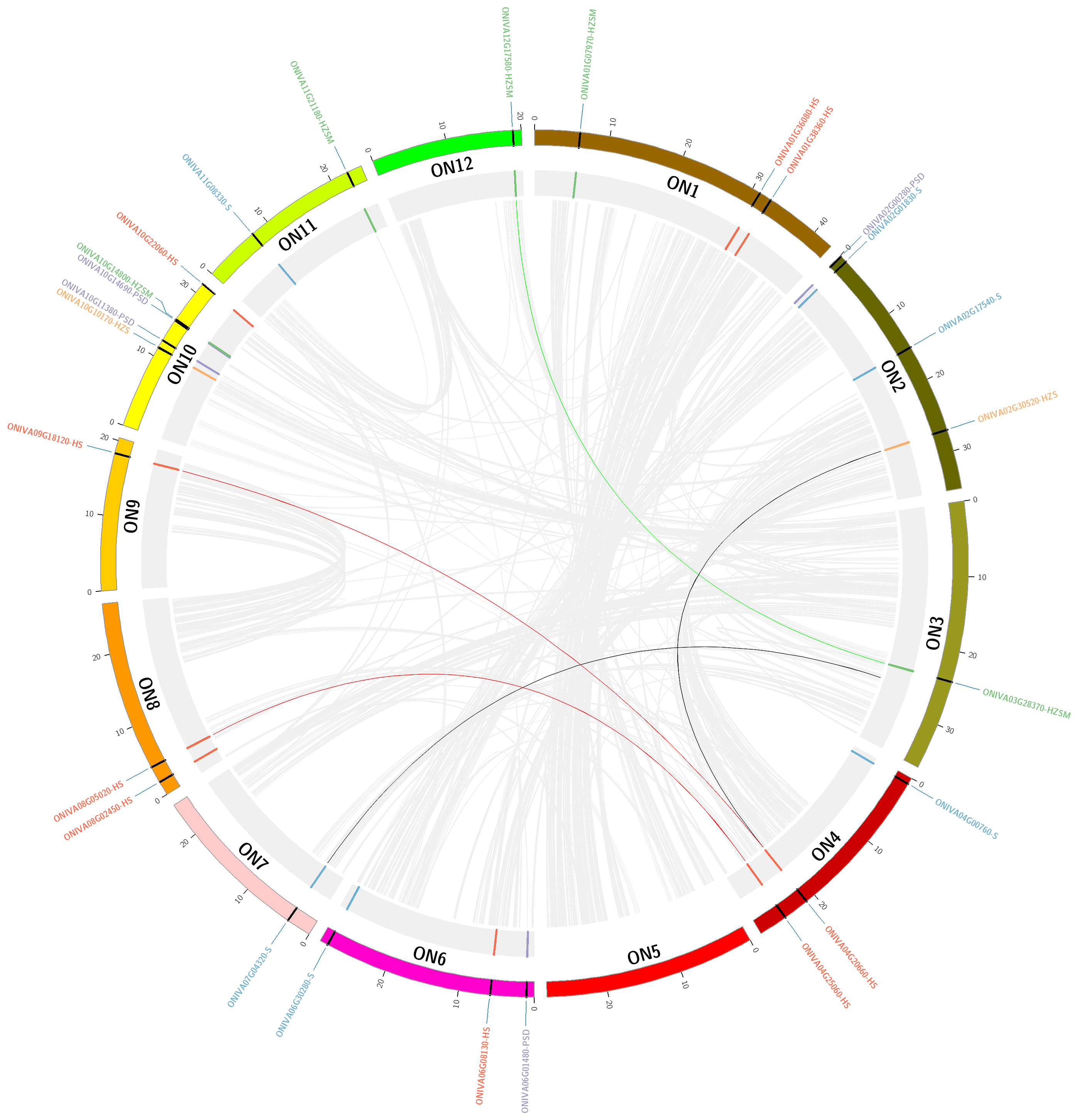

**Supplementary Figure S2E.** Collinear blocks of the *Oryza nivara* genome.

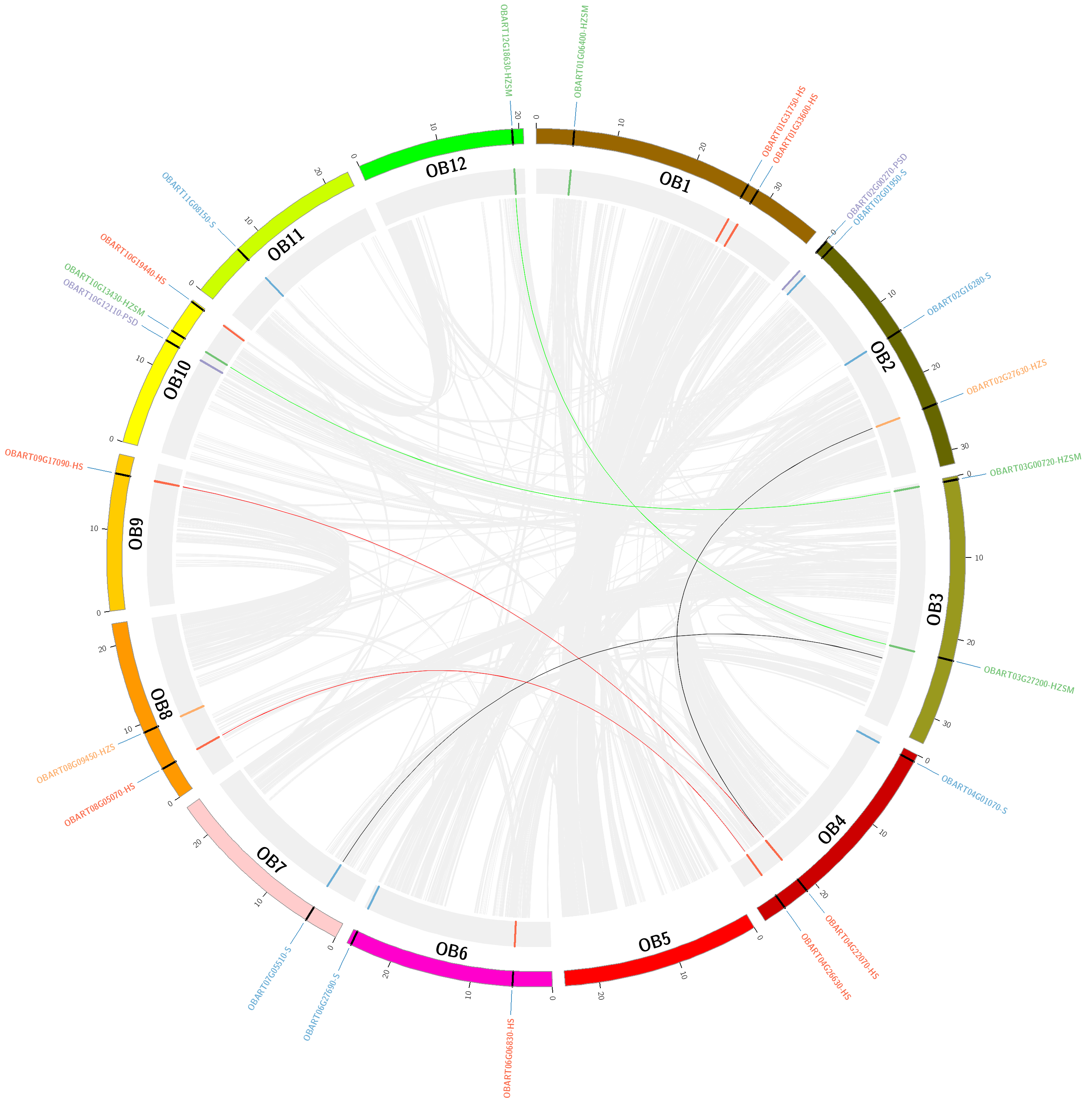

**Supplementary Figure S2F.** Collinear blocks of the *Oryza barthii* genome

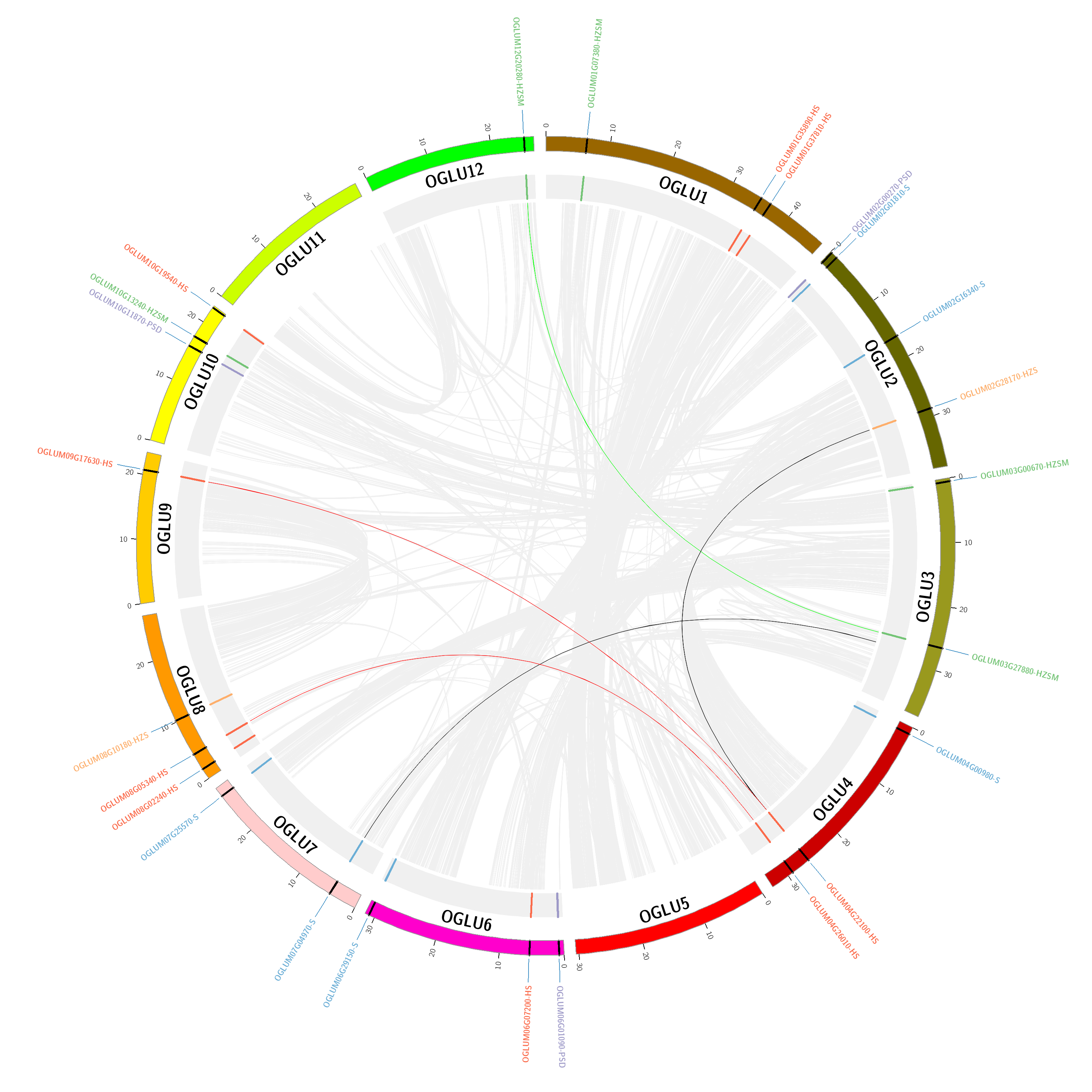

**Supplementary Figure S2G.** Collinear blocks of the *Oryza glumaepatula* genome.

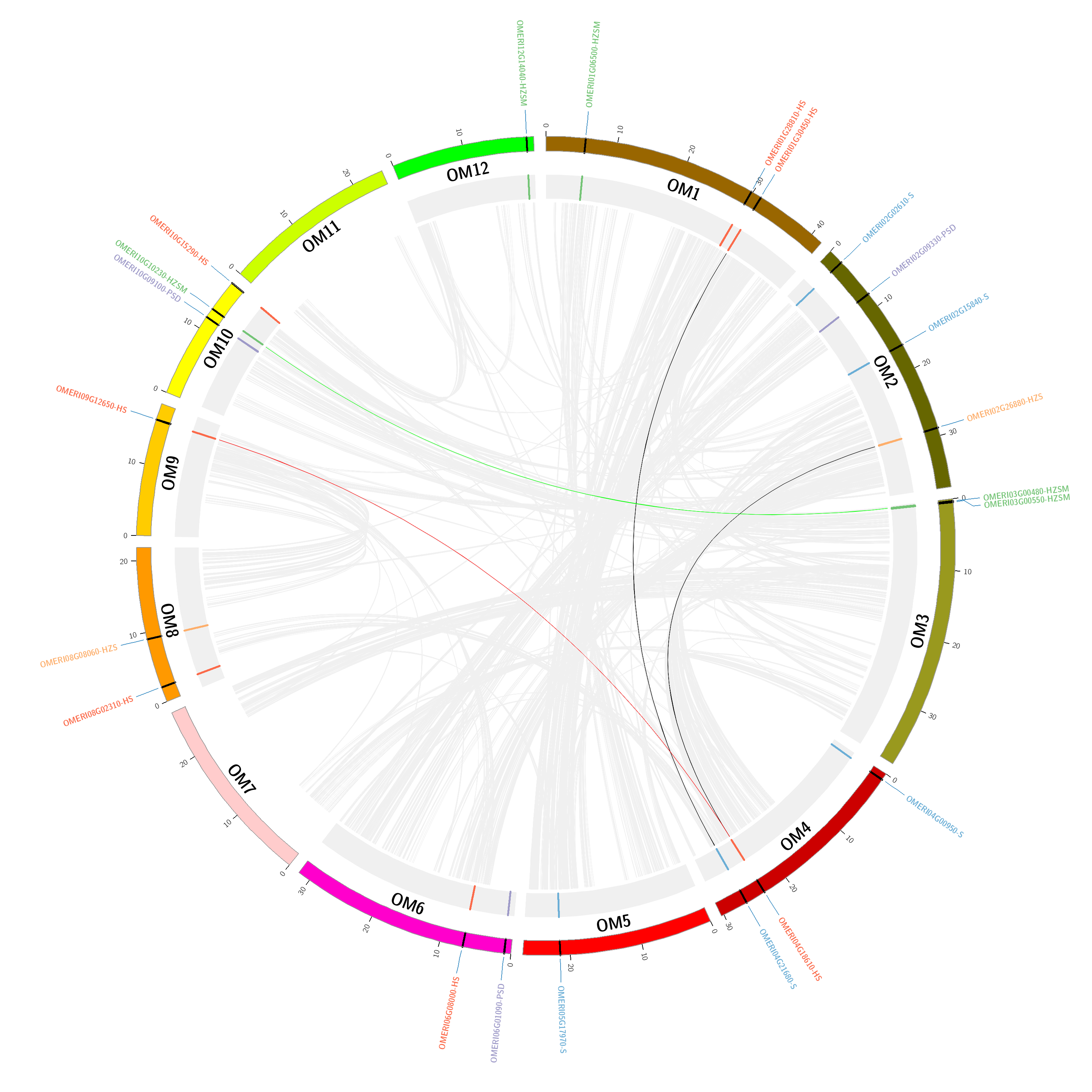

**Supplementary Figure S2H.** Collinear blocks of the *Oryza meridionalis* genome.

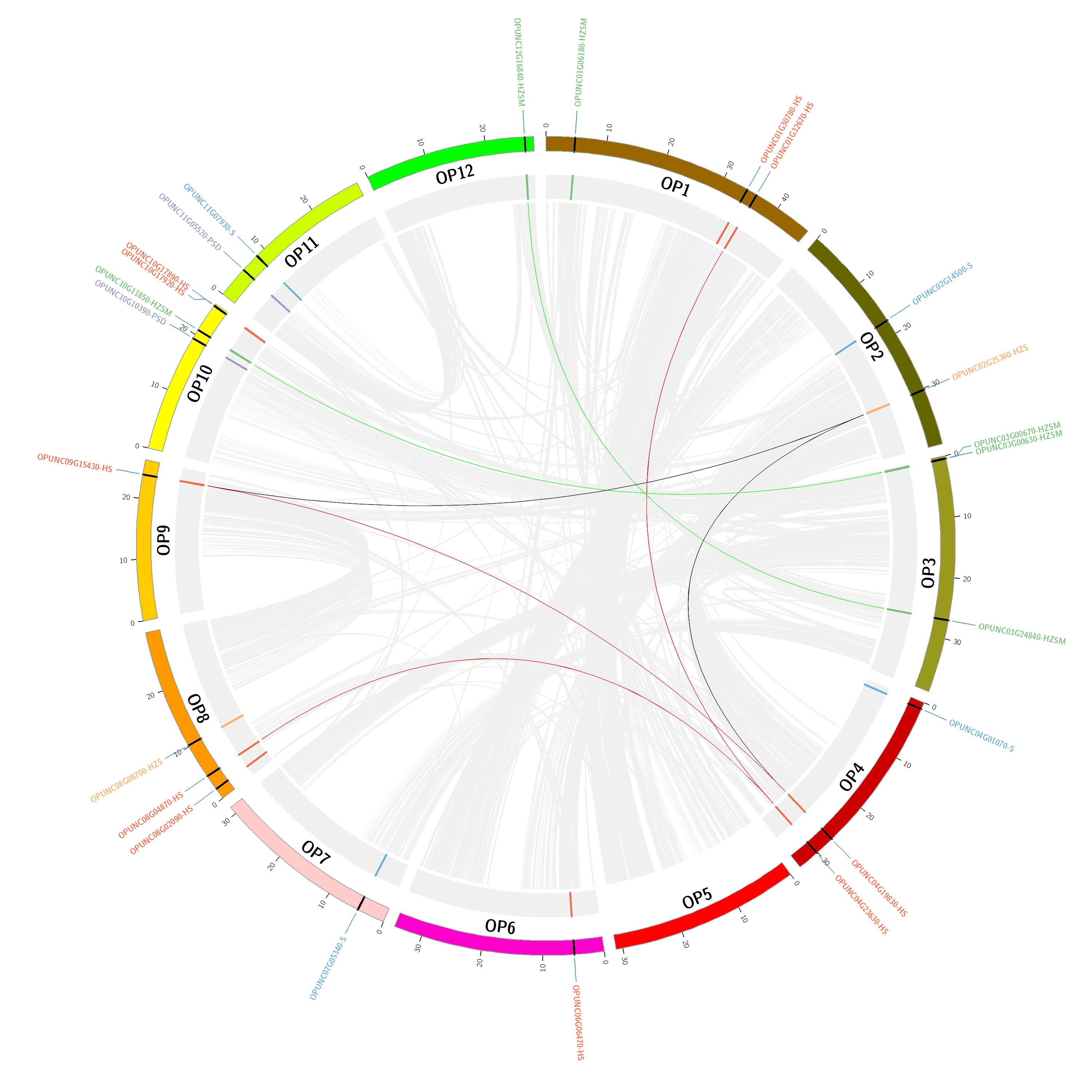

**Supplementary Figure S2I.** Collinear blocks of the *Oryza punctata* genome.

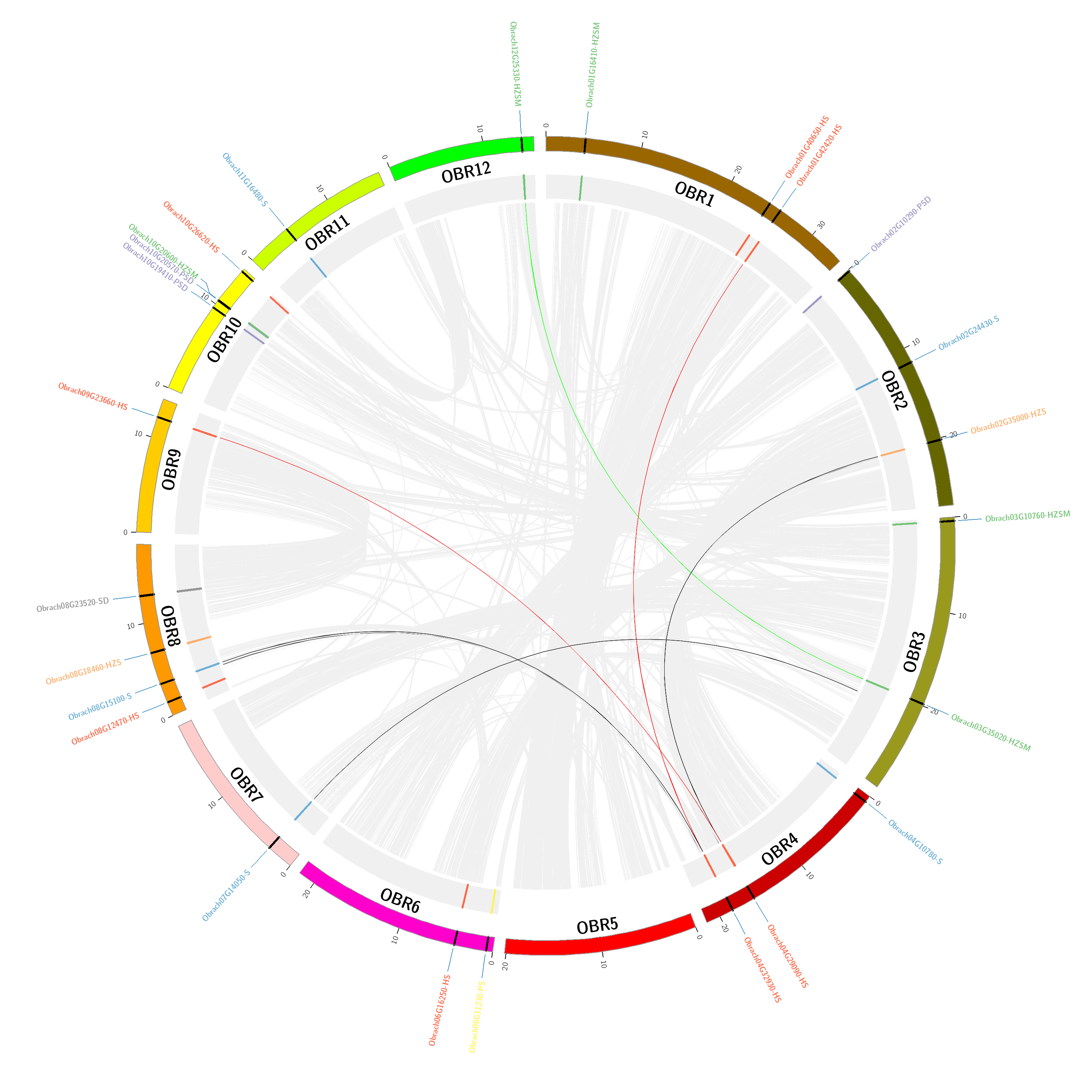

**Supplementary Figure S2J.** Collinear blocks of the *Oryza brachyantha* genome.

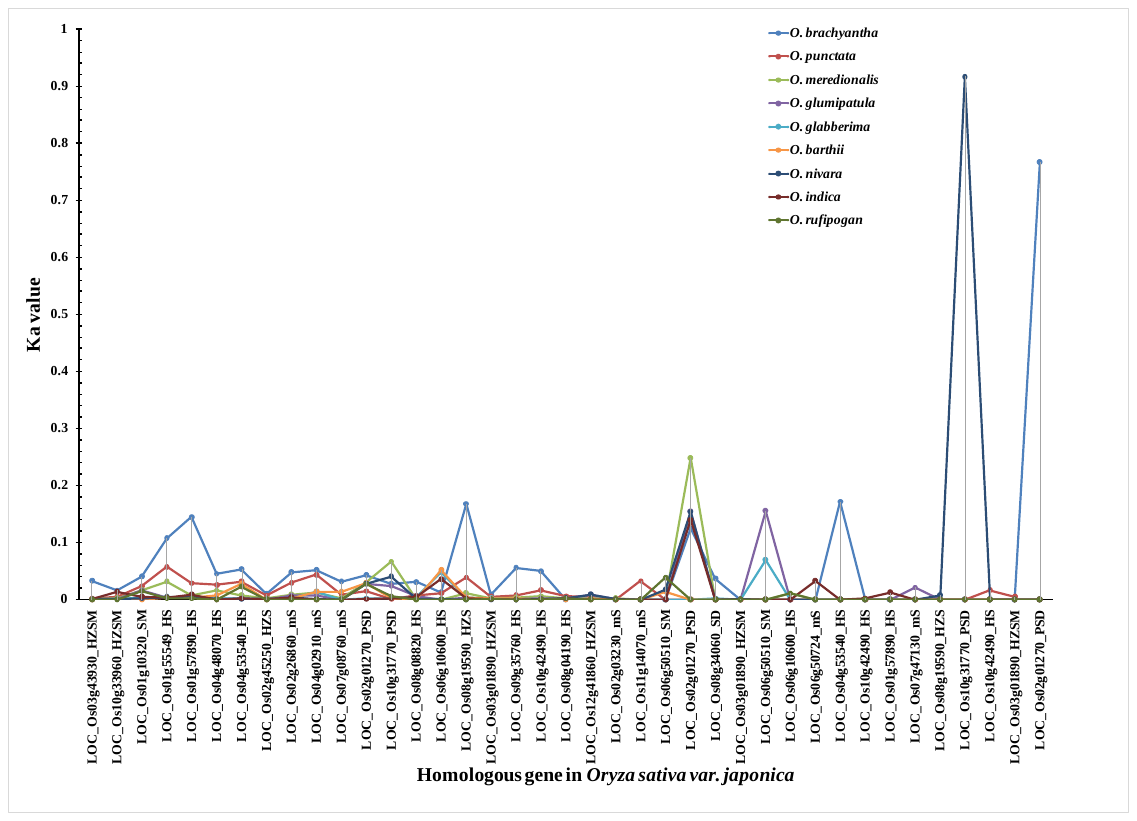

**Supplementary Figure S3A.** Ka values for START homologues of different Oryza species with respect to *Oryza sativa var. japonica*

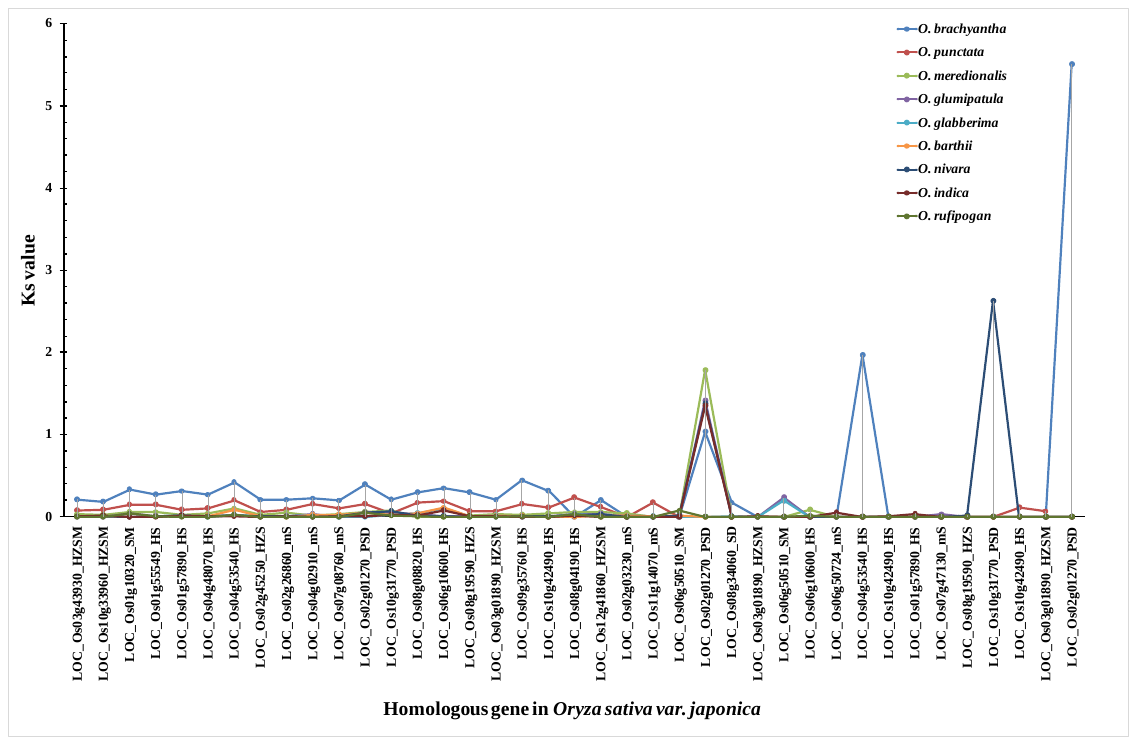

**Supplementary Figure S3B.** ks values for START homologues of different Oryza species with respect to *Oryza sativa var. japonica*

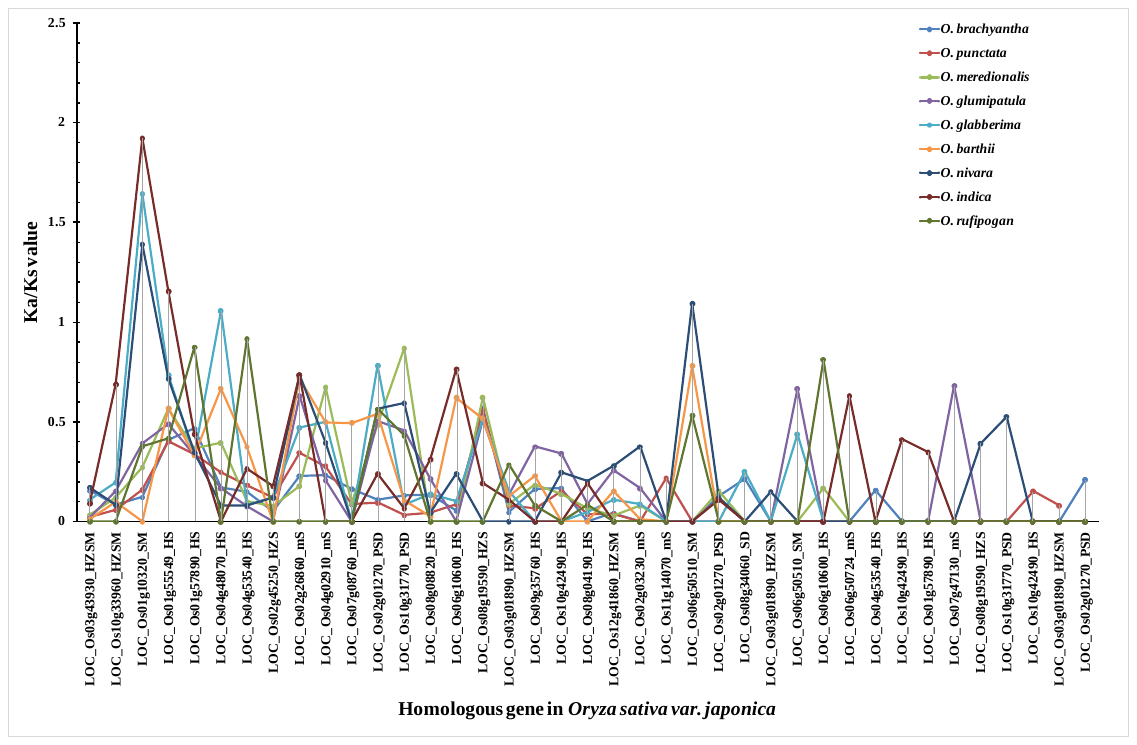

**Supplementary Figure S3C.** Ka/ks values for START homologues of different Oryza species with respect to *Oryza sativa var. japonica*
