## Supplementary Table S3 for "Comparative genomics of StAR-related lipid transfer (START) domains across wild and cultivated rice"

**Supplementary Table S3.** Collinear genes number across ten Oryza genome.

| **Name of Oryza Species****`** | **Total number of genes** | **Number of collinear genes** | **Percentage of Collinear genes** | **Number of START genes in Collinear blocks** |
| --- | --- | --- | --- | --- |
| **Cultivated Rice Species** | | | | |
| *Oryza sativa var. japonica* | 42189 | 6505 (200) | 15.42 | 10 |
| *Oryza indica var. indica* | 42031 | 6116 (171) | 14.55 | 6 |
| *Oryza glaberrima* | 34130 | 6376 (255) | 18.68 | 7 |
| **Wild Rice Species** | | | | |
| *Oryza rufipogon* | 37912 | 6057 (176) | 15.98 | 8 |
| *Oryza nivara* | 37026 | 4559 (180) | 12.31 | 8 |
| *Oryza barthii* | 35553 | 5622 (166) | 15.81 | 10 |
| *Oryza glumaepatula* | 36379 | 5927 (171) | 16.29 | 8 |
| *Oryza meridionalis* | 30241 | 3944 (151) | 13.04 | 7 |
| *Oryza punctata* | 32550 | 6216 (174) | 19.10 | 10 |
| *Oryza brachyantha* | 32463 | 5723 (154) | 17.63 | 9 |

(#) Number of alignment blocks across the genome is mentioned in parentheses
