## Supplementary Table S4 for "Comparative genomics of StAR-related lipid transfer (START) domains across wild and cultivated rice"

**Supplementary Table S4.** Ka Ks, and Ka/Ks analysis among the gene pairs that follow different modes of duplication in *Oryza sativa var japonica* genome.

| Gene pair | Mode of duplication | Ka | Ks | Ka/Ks |
| --- | --- | --- | --- | --- |
| LOC_Os06g50560_mS-LOC_Os06g50510_SM  SM is having higher expression levels while the mS is having least or below basal level of expression claiming its loss of gene function. Similar is the case in the anatomical parts development supporting loss of function. | Proximal | 0.052 | 0.069 | 0.762 |
| LOC_Os06g10600_HS-LOC_Os07g47130_mS  HS showed higher stage specific expression in the developmental stages and the newly transposed pair mS showed expression in the same stages of its parent gene but very low levels also indicates loss of function or importance.  Similar is the case in the anatomical parts development. | Transposed | 0.446 | 2.016 | 0.365 |
| LOC_Os02g26860_mS-LOC_Os04g02910_mS  Parental mS showed stage specific expression the derivative also showed similar stage specific expression but an above optimal levels which indicates the stage specific memory of the parental influence and additionally it showed expression in some other stages where parent gene has not showed expression indicating its unassigned role in developmental stage specificity. Similar is the case in anatomical parts development. | Transposed | 1.044 | 6.057 | 0.392 |
| LOC_Os10g33960_HZSM-LOC_Os03g01890_HZSM  Dev: (similar trend but slight variation in the expression level)  Organellar/anatomical parts development:  Almost same | Segmental | 0.062 | 1.017 | 0.063 |
| LOC_Os12g41860_HZSM-LOC_Os03g43930_HZSM  Dev: (no similar trend except equal level of expression in isolated stages)  Organellar:  Almost same | Segmental | 0.081 | 0.642 | 0.127 |
| LOC_Os08g08820_HS-LOC_Os04g53540_HS  Dev: (almost same level of expression with similar trend)  Functional siginificance in anatomical parts too with mirror image type of stage specific expression levels. | Segmental | 0.086 | 1.070 | 0.117 |
| LOC_Os04g48070_HS-LOC_Os02g45250_HZS  Dev: (almost similar trend and similar level of expression except slight variation in the stage level expression)  Loss of bzip fragment may have affected its expression in certain stages when compared to the HZS partner in anatomical parts development. | Segmental | 0.177 | 1.134 | 0.166 |
| LOC_Os04g48070_HS-LOC_Os09g35760_HS  Dev: (almost similar trend and similar level of expression except slight variation in the stage level expression).  HS on chromosome 9 had higher expression in almost all stages indicating its functional significance. | Segmental | 0.341 | 2.923 | 0.160 |
